# Identifying molecular targets of a colloidal nanosilver formulation (Silversol^®^) in multidrug resistant *Pseudomonas aeruginosa*

**DOI:** 10.1101/2023.06.27.546803

**Authors:** Gemini Gajera, Nidhi Thakkar, Chhaya Godse, Anselm DeSouza, Dilip Mehta, Vijay Kothari

## Abstract

*P. aeruginosa* is a notorious pathogen. A multi-drug resistant strain of this bacterium was challenged with a colloidal nano-silver formulation- Silversol^®^. Its minimum inhibitory concentration against *P. aeruginosa* was found to be 1.5 ppm, and at sub-MIC of 1 ppm, it was able to alter quorum-sensing regulated pigmentation, exopolysaccharide synthesis and biofilm formation, antibiotic susceptibility, protein synthesis and export, nitrogen metabolism, and siderophore production in this pathogen. Transcriptome analysis of the silver-exposed *P. aeruginosa* indicated generation of nitrosative stress and disturbance of iron homeostasis to be the major mechanisms associated with anti-Pseudomonas activity of Silversol^®^. Network analysis of the differentially expressed genes in silver-treated bacterium identified ten genes as the potential molecular targets: norB, norD, nirS, nirF, nirM, nirQ, nosZ, nosY, narK1, and norE (all associated with nitrogen metabolism or denitrification). Three of them (norB, narK1, and norE) were also validated through RT-PCR.

## 1. Introduction

Silver has a long history of therapeutic use in traditional medicine. In the modern era too, many reports (Zhang et al., 2016; 2. Anandaradje et al., 2020; Wahab et al., 2021) have been published describing various biological activities of silver. Antimicrobial activity of different forms of silver are also well known e.g., antibacterial (Tang et al., 2018; Balachandar et al., 2022), antifungal (Matras et al., 2022), antiviral (Salleh et al., 2020), antiprotozoal (Hassan et al., 2019), and anthelmintic (Gajera et al., 2023). Different forms of silver (e.g., metallic, colloidal, silver salts) may have different modes of action, and varying degree of biological activity. Antibacterial activity of silver becomes even more important in face of the resistance displayed by bacterial pathogens against conventional antibiotics. Till now, there is not much indication from literature about bacteria developing resistance against silver. In general, silver is believed to be killing bacteria by altering cell membrane permeability, generating reactive oxygen species (ROS), and interrupting DNA replication (Ahmad et al., 2017; Mazur et al., 2020; Reeves et al., 2020; Yin et al., 2020; Jiang et al., 2022). However, much investigation still is required to elucidate precise molecular mechanisms associated with antibacterial activity of silver.

Since metallic form of silver is known to be more potent than its ionic form (de Souza et al., 2021), we chose a solution of colloidal silver for the purpose of this study. Target pathogen for this study was *Pseudomonas aeruginosa*, one of the most notorious and versatile bacterial pathogens. Its antibiotic-resistant strains have been listed by Centers for Disease Control and Prevention (CDC; https://www.cdc.gov/drugresistance/biggest-threats.html), World Health Organization (WHO; https://www.who.int/publications/i/item/WHO-EMP-IAU-2017.12), and Department of Biotechnology of the Indian government (DBT; https://dbtindia.gov.in/sites/default/files/IPPL_final.pdf) among priority pathogens, against whom there is an urgent need to discover new antibiotics (Weiner et al., 2016). This study investigated effect of Silversol^®^ (a colloidal nano-silver formulation) on *P. aeruginosa*’s growth, various virulence traits, and gene expression profile at the whole transcriptome level.

## 2. Methods

### 2.1. Colloidal silver formulation

The test formulation Silversol^®^ (32 ppm) originally developed by American Biotech Labs (USA) was procured from Viridis BioPharma Pvt Ltd, Mumbai, India. It is a colloidal silver preparation reported to possess multiple biological activities (de Souza et al., 2021). The elemental form of zero-valent metallic silver particles contained in this product is coated with silver oxide, and the particle size is claimed by the manufacturer to range between 5-50 nm.

### 2.2. Bacterial culture

The *P. aeruginosa* strain used in this study was sourced from our internal culture collection. This strain has been well characterized by us with respect to its antibiotic resistance, pigment production and certain other virulence traits. Its antibiogram generated through a disc-diffusion assay performed as per NCCLS guidelines revealed it to be resistant to 8 antibiotics (co-trimoxazole, augmentin, nitrofurantoin, ampicillin, chloramphenicol, clindamycin, cefixime, and vancomycin) belonging to 5 different classes (Fluoroquinolones, Beta lactams, Third generation cephalosporins, Macrolides, and Sulfonamides). Hence it can be described as a multidrug resistant (MDR) strain. As reported in our earlier publications (Joshi et al., 2019; Patel et al., 2019) involving this strain, it is a haemolytic strain capable of producing the quorum sensing (QS) regulated pigments (pyocyanin and pyoverdine), and also of biofilm formation.

As we also had *P. aeruginosa* PAO1 (MTCC 3541) strain in our lab, we could compare the MDR strain used in this study with the PAO1 in terms of their antibiotic resistance and virulence towards *C. elegans*. The strain used in our experiments was resistant to more classes of antibiotics, and it also displayed higher virulence towards the model host *Caenorhabditis elegans* (Table S1; Figure S1).

This bacterium was maintained on Pseudomonas agar (HiMedia). While culturing the bacteria for *in vivo* assay, they were grown in Pseudomonas broth (magnesium chloride 1.4 g/L, potassium sulphate 10 g/L, peptic digest of animal tissue 20 g/L, pH 7.0 ± 0.2) supplemented with glycerol (3%v/v; HiMedia).

### 2.3. Quantification of growth and quorum-regulated pigments

Effect of Silversol^®^ on bacterial growth and pigment formation was quantified through broth dilution assay. Following incubation of *P. aeruginosa* in Pseudomonas broth supplemented with or without Silversol^®^ for 21±1 h at 35°C, cell density was measured at 764 nm (Agilent Cary 60 UV-Vis). Pigments from the culture broth were extracted as described in Unni et al., (2014) and El-Fouly et al., (2015). One mL of culture broth was mixed in a 2:1 ratio with chloroform (Merck, Mumbai), followed by centrifugation (15,300 g) for 10 min. This resulted in formation of two immiscible layers. OD of the upper aqueous layer containing the yellow-green fluorescent pigment pyoverdine was measured at 405 nm. Pyoverdine Unit was calculated as OD_405_/OD_764_. The lower chloroform layer containing the blue pigment pyocyanin was mixed with 0.1 N HCL (20%v/v; Merck). This caused a change of colour from blue to pink. This was followed by centrifugation (15,300 g) for 10 min, and OD of upper layer acidic liquid containing pyocyanin was quantified at 520 nm. Pyocyanin Unit was calculated as OD_520_/OD_764._

The lowest concentration of silver capable of inhibiting ≥80% growth was taken as minimum inhibitory concentration (MIC) (Turnidge, 2015). From each tube showing absence of growth, 0.1 mL of broth was spread on Pseudomonas agar, and these agar plates were observed for appearance of growth over an incubation (at 35°C) period of 72 h for determination of minimum bactericidal concentration (MBC). Extended incubation till 72 h was made in this case to be able to differentiate true bactericidal effect from any possible post-antibiotic effect (Ramadan et al., 1995; Pfaller et al., 2004). The concentration of silver which could inhibit appearance of growth on agar plates completely was taken as MBC.

### 2.4. Haemolysis assay

Haemolytic activity is considered to be an important virulence trait of pathogens including *P. aeruginosa*, particularly under iron-limiting conditions. To investigate whether silver-exposure can have any impact on haemolytic potential of *P. aeruginosa*, we inoculated the silver-pre-treated cells on blood agar plate (HiMedia) and compared the haemolysis pattern with that of control.

OD_764_ of *P*. *aeruginosa* grown in Pseudomonas broth in presence or absence of Silversol^®^ was adjusted to 1.0, and 20 µL of this culture suspension was added onto the center of blood agar plate, followed by incubation at 35°C for 24 h. Next day plates were observed for a zone of haemolysis surrounding the point of inoculation.

### 2.5. Antibiotic susceptibility assay

Antibiogram of *P. aeruginosa*’s overnight grown culture in Pseudomonas broth in presence or absence of Silversol^®^ was generated through disc diffusion assay in accordance to NCCLS guidelines (Lacy et al., 2004). Cells grown in Pseudomonas broth were separated through centrifugation (13600 g) and washed with phosphate buffer (pH 7.0±0.2) followed by centrifugation. The resulting cell pellet was used to prepare inoculum for subsequent disc diffusion assay by suspending the cells in normal saline and adjusting the OD_625_ between 0.08-0.10 to achieve an inoculum density equivalent to McFarland standard 0.5. One hundred µL of this inoculum was spread onto cation-adjusted Muller-Hinton agar (HiMedia) plates (Borosil; 150 mm) followed by placing the antibiotic discs (Icosa G-I MINUS; HiMedia, Mumbai) on the agar surface. Incubation at 35°C was made for 18±1 h, followed by observation and measurement of zone of inhibition.

### 2.6. Biofilm assay

Biofilm formation is an important virulence trait, and hence effect of Silversol^®^ on biofilm forming ability of *P. aeruginosa*, as well as on pre-formed biofilm was investigated. In this assay, control and experimental, both groups contained nine test tubes (15 mL). In each group, three subgroups were made. First subgroup of three test tubes in the experimental group contained Pseudomonas broth supplemented with Silversol^®^ (1 ppm), whereas remaining six tubes contained Pseudomonas broth with no Silversol^®^ on first day of experiment. All these tubes were inoculated with inoculum (10% v/v) standardized to 0.5 McFarland turbidity standard (making total volume in tube 1 mL), followed by incubation at 35°C for 24 h under static condition, which resulted in formation of biofilm as a ring on walls of the glass tubes., This biofilm was quantified by crystal violet assay (Hirshfield et al., 2009), preceded by quantification of bacterial cell density and pigment. Content from the remaining six test tubes from rest of the two subgroups were discarded following cell density and pigment estimation, and then the biofilms remaining on inner surface of these tubes were washed with phosphate buffer saline (PBS; pH: 7±0.2) to remove loosely attached cells. Now, 2 mL of minimal media (Sucrose 15 g/L, K_2_HPO_4_ 5.0 g/L, NH_4_Cl_2_ g/L, NaCl 1 g/L, MgSO_4_ 0.1 g/L, yeast extract 0.1 g/L, pH 7.4±0.2) containing Silversol^®^ (1 ppm), was added into each of these tubes, so as to cover the biofilm completely, and tubes incubated for 24 h at 35°C. At the end of incubation, one subgroup of 3 tubes was subjected to crystal violet assay to know whether any eradication of the pre-formed biofilm has occurred under the influence of Silversol^®^, and the last subgroup of 3 tubes was subjected to viability assessment through MTT assay (Trafny et al., 2013).

For the crystal violet assay, the biofilm containing tubes (after discarding the inside liquid) were washed with PBS in order to remove all non-adherent (planktonic) bacteria, and air-dried for 15 min. Then, each of the washed tubes was stained with 1.5 mL of 0.4% aqueous crystal violet (Central Drug House, Delhi) solution for 30 min. Afterwards, each tube was washed twice with 2 mL of sterile distilled water and immediately de-stained with 1.5 mL of 95% ethanol. After 45 min of de-staining, 1 mL of de-staining solution was transferred into separate tubes, and read at 580 nm (Agilent Cary 60 UV-Vis).

For the MTT assay, the biofilm-containing tubes (after discarding the inside liquid) were washed with PBS in order to remove all non-adherent (planktonic) bacteria, and air-dried for 15 min. Then 900 μL of minimal media was added into each tube, followed by addition of 100 μL of 0.3% MTT [3-(4,5-Dimethylthiazol-2-yl)-2,5-iphenyltetrazolium Bromide; HiMedia]. Then after 2 h incubation at 35°C, all liquid content was discarded, and the remaining purple formazan derivatives were dissolved in 2 mL of DMSO and measured at 540 nm.

### 2.7. Exopolysaccharide (EPS) quantification

*P. aeruginosa* was grown in 100 mL flasks containing 20 mL of Pseudomonas broth. Incubation at 35°C was made for 24 h with intermittent shaking. Following estimation of growth by measuring OD at 764 nm, culture broth was subjected to centrifugation (13,600 g for 10 min), and the supernatant was used for EPS quantification using the method described in Li et al. (2012) with some modification. Briefly, 40 mL of chilled acetone (Merck) was added to 20 mL of supernatant, and allowed to stand for 30 min. The EPS precipitated thus was separated by filtration through pre-weighed Whatman # 1 filter paper (Whatman International Ltd., England). Filter paper was dried at 60° C for 24 h, and weight of EPS on paper was calculated.

### 2.8. Protein estimation

Extracellular protein present in bacterial culture (grown in presence or absence of silver) supernatant, and intracellular protein in the cell lysate was quantified through Folin-Lowry method (Lowry et al., 1951; Dulley and Grieve et al., 1975). After measuring cell density, 1 mL of *P. aeruginosa* culture grown in Pseudomonas broth was centrifuged (13,600 g), and the resulting supernatant was used for extracellular protein estimation. The remaining cell pellet was subjected to lysis (Mishra et al., 2017) for release of intracellular proteins. Briefly, the cell pellet was washed with phosphate buffer (pH 7.4), and centrifuged (13,600 g). Resulting pellet was resuspended in 1 mL of chilled lysis buffer (8.76 g/L NaCl, 10 mL of Triton X 100, 5/L g sodium deoxycholate, 1 g/L sodium dodecyl sulphate, and 6 g/L Tris HCl, in 990 mL of distilled water), and centrifuged (500 rpm) for 30 min at 4°C for agitation purpose. This was followed by further centrifugation (16,000 g at 4°C) for 20 min. Resulting cell lysate (supernatant) was used for protein estimation. Kanamycin (at IC_50_: 200 µg/mL), an aminoglycoside antibiotic known to inhibit bacterial protein synthesis (Suzuki et al., 1970), was used as a positive control.

### 2.9. Nitrite estimation

Quantification of nitrite in bacterial culture was achieved through a colorimetric assay using modified Griess reagent (Misko et al., 1993). A total of 250 µL of supernatant obtained from centrifugation of *P. aeruginosa* culture grown in presence or absence of silver, subjected to centrifugation at 13,500 g for 10 minutes at 25°C, was mixed with an equal volume of Griess reagent (1X concentration; Sigma-Aldrich) followed by 15 min incubation in dark at room temperature. Absorbance of the resulting colour was measured at 540 nm. This absorbance was plotted on the standard curve prepared using NaNO_2_ (0.43-65 µM) to calculate nitrite concentration. Sodium nitroprusside (Sigma Aldrich) was used as a positive control, as it is known to generate nitrosative stress in bacteria (Auger et al., 2011).

### 2.10. Transcriptome analysis

To gain insights into the molecular mechanisms through which silver could inhibit bacterial growth and modulate various traits like QS, gene expression profile of *P. aeruginosa* challenged with sub-MIC of Silversol^®^ (1 ppm) was compared with that of control culture at the whole transcriptome level. Overall workflow of this whole transcriptome analysis (WTA) aimed at obtaining a holistic picture regarding mode of action of this formulation is presented in Figure S2.

#### 2.10.1. RNA extraction

RNA from bacterial cells was extracted by Trizol (Invitrogen Bioservices; 343909) method. Precipitation was done by isopropanol followed by washing with 75% ethanol, and RNA was dissolved in nuclease free water. Extracted RNA was quantified on Qubit 4.0 fluorometer (Thermofisher; Q33238) using RNA HS assay kit (Thermofisher; Q32851) following manufacturer’s protocol. Purity and concentration of RNA was measured on Nanodrop 1000. Finally, to obtain RIN (RNA Integrity Number) values, RNA was checked on the TapeStation using HS RNA ScreenTape (Agilent) (Table S2).

#### 2.10.2. Library preparation

Final libraries were quantified through Qubit 4.0 fluorometer (Thermofisher; Q33238) using DNA HS assay kit (Thermofisher; Q32851) following manufacturer’s protocol. To identify the insert size of the library, it was queried on Tapestation 4150 (Agilent) employing high sensitive D1000 screentapes (Agilent; 5067-5582) as per manufacturers’ protocol. Acquired sizes of all libraries are reported in Table S3.

#### 2.10.3. Genome annotation and functional analysis

Quality assessment of the raw fastq reads of the sample was performed using FastQC v.0.11.9 (default parameters) (Andrews, 2010). The raw fastq reads were pre-processed using Fastp v.0.20.1 (Chen et al., 2018), followed by quality reassessment using FastQC.

The *P. aeruginosa* genome (GCA_000006765.1_ASM676v1) was indexed using bowtie2-build (Langmead and Salzberg, 2012) v2.4.2 (default parameters). The processed reads were mapped to the indexed *P. aeruginosa* genome using bowtie2 v2.4.2 parameters. The aligned reads from the individual samples were quantified using feature count v. 0.46. 1 (Liao et al., 2014) to obtain gene counts. These gene counts were used as inputs to edgeR (Robinson et al., 2010) using exact test (parameters: dispersion 0.1) for differential expression estimation. The up- and down-regulated sequences were extracted from the *P. aeruginosa* coding file and subjected to blast2go (Conesa et al., 2008) for annotation to extract the Gene Ontology (GO) terms. These GO terms were subjected to the wego (Ye et al., 2018) tool to obtain gene ontology bar plots.

All the raw sequence data has been submitted to Sequence Read Archive. Relevant accessions no. is SRX14392191 (https://www.ncbi.nlm.nih.gov/sra/SRX14392191).

### 2.11. Network analysis

From the list of differentially expressed genes (DEG) in Silversol^®^-exposed *P. aeruginosa*, those satisfying dual filter criteria of log fold-change ≥2 and FDR ≤0.05 were chosen for further network analysis. The list of such DEGs was fed into the STRING (v.11.5) database (Szklarczyk et al., 2019) to generate the PPI (Protein-Protein Interaction) network. Members of this PPI network were then arranged in decreasing order of ’node degree’ (a measure of connectivity with other genes or proteins), and those above a specified threshold value were subjected to ranking by the cytoHubba plugin (v.3.9.1) (Chin et al., 2014) of Cytoscape (Shannon et al., 2003). As cytoHubba employs 12 different ranking methods, we considered the DEG being top-ranked by ≥6 different methods (i.e., 50% of the total ranking methods) for further investigation. These top-ranked shortlisted proteins were then subjected to local cluster analysis through STRING, and those that were part of multiple clusters were termed potential ’hubs’ that can be investigated for additional validation of their targetability. The term ’hub’ refers to a gene or protein that interacts with multiple other genes/proteins. The identified hubs were then subjected to co-occurrence analysis to see whether an antibacterial agent targeting them is likely to meet the requirement of selective toxicity (targeting the pathogen while causing no harm to the host). This sequence of analysis allowed us to end up with a limited number of proteins that satisfied multiple statistical and biological significance criteria simultaneously: (i) log fold-change ≥2; (ii) FDR≤0.05; (iii) relatively higher node degree; (iv) top-ranking by at least 6 cytoHubba methods; (v) (preferably) member of more than 1 local network cluster; and (vi) high probability of the target being absent from the host.

### 2.12. Polymerase Chain Reaction (RT-PCR)

Differential expression of the potential hubs identified through network analysis of DEG revealed from WTA was confirmed through PCR too. Primer designing for the selected genes was accomplished through Primer3 Plus (Untergasser et al., 2007). Primer sequences thus obtained were confirmed for their binding withing the whole *P. aeruginosa* genome exclusively to the target gene sequence. Primer sequences for all the target genes are listed in Table 1. RNA extraction and quality check was done as described in preceding section. cDNA was obtained by using SuperScript™ VILO™ cDNA Synthesis Kit (Invitrogen Biosciences). PCR assay was conducted by using gene specific primers procured from Sigma-Aldrich. The gene PA3617 (recA) was kept as an endogenous control. The reaction mix used was FastStart Essential DNA Green Master mix (Roche; 06402712001). Real time PCR assay was performed on Quant studio 5 real time PCR machine (Thermo Fisher Scientific, USA). Temperature profile employed is provided in Table S4.

**Table 1.**
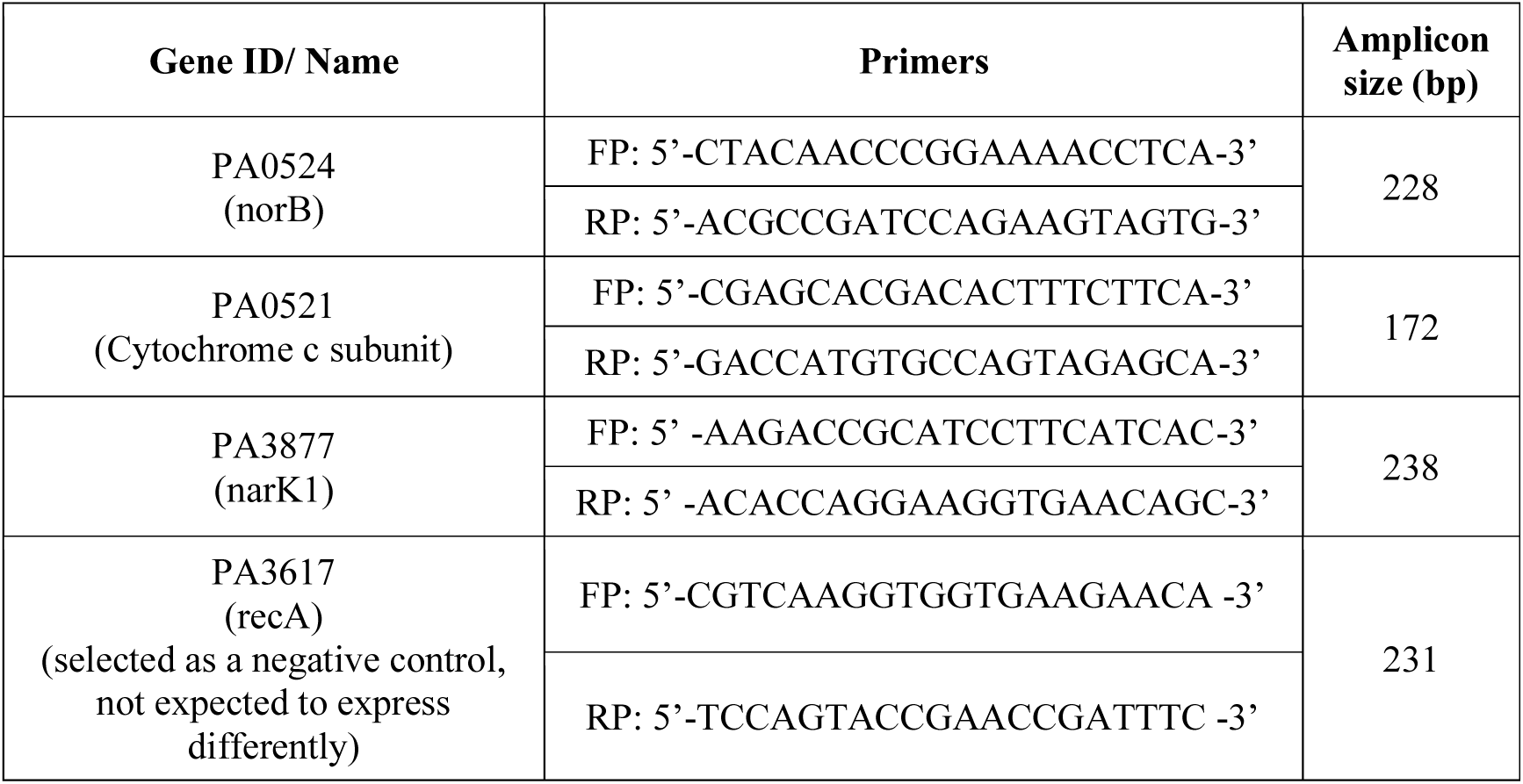
Primer sequences for the target genes.

**Table 2.**
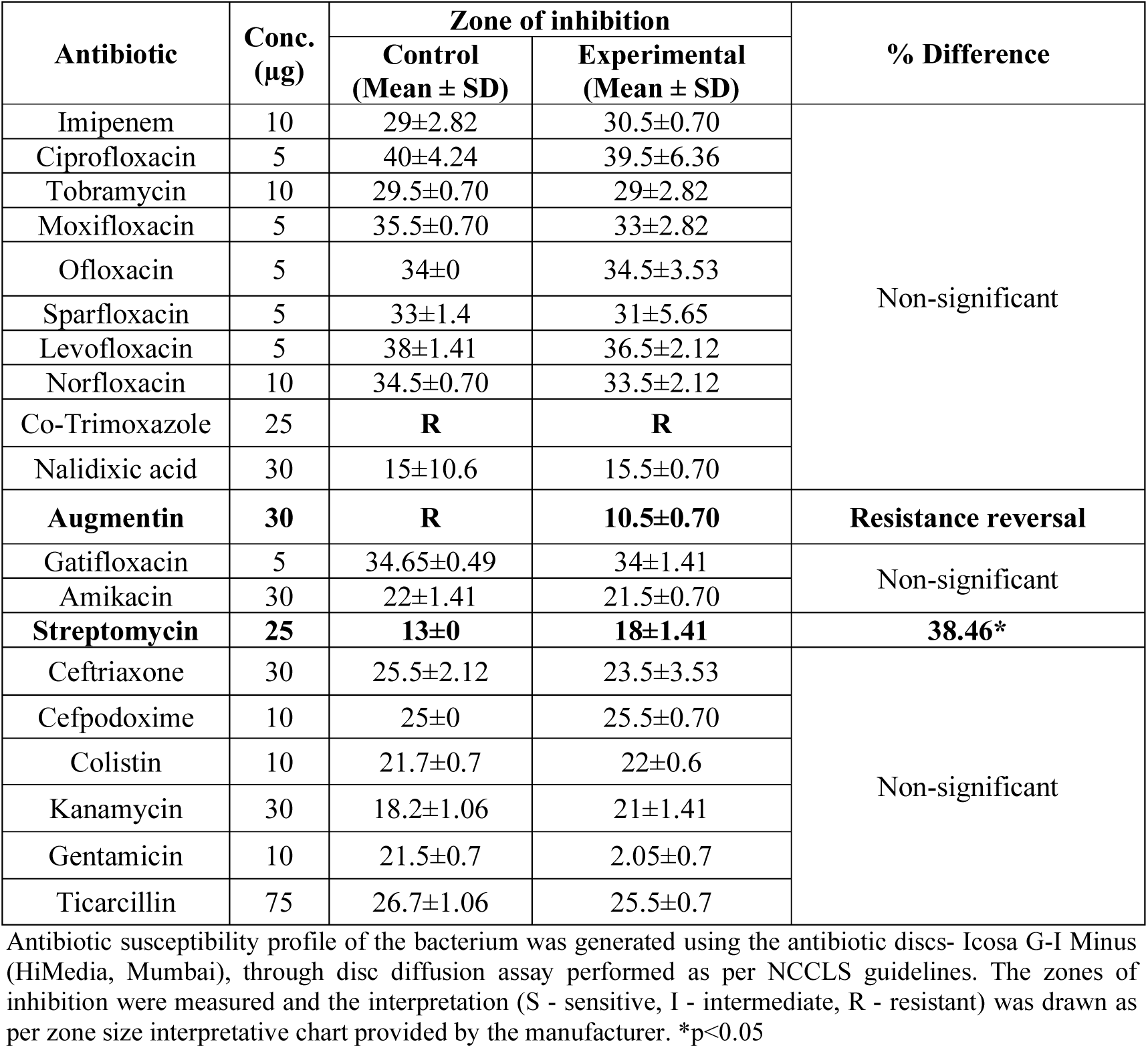
Silver pre-treatment modulates bacterial susceptibility to some antibiotics.

**Table 3.** List of DEGs in Silversol^®^ exposed *P. aeruginosa* satisfying the dual criteria of log fold-change ≥2 and FDR≤0.05.

| Sr. No. | Gene ID | Gene Symbol | Codes for | Log FC | FDR | Up- or down-regulation |
| --- | --- | --- | --- | --- | --- | --- |
| 1 | PA0522 | NA | hypothetical protein | 4.9 | 5.87E-06 | ↑ |
| 2 | PA2691 | NA | conserved hypothetical protein | 4.2 | 5.10284E-05 | ↑ |
| 3 | PA2689 | NA | hypothetical protein | 4.02 | 0.0002 | ↑ |
| 4 | PA0521 | NA | probable cytochrome c oxidase subunit | 3.91 | 0.0002 | ↑ |
| 5 | PA4623 | NA | hypothetical protein | 3.5 | 0.001 | ↓ |
| 6 | PA3237 | NA | hypothetical protein | 3.47 | 0.001 | ↓ |
| 7 | PA0524 | norB | nitric-oxide reductase subunit B | 3.41 | 0.001 | ↑ |
| 8 | PA3877 | narK1 | nitrite extrusion protein 1 | 3.27 | 0.003 | ↑ |
| 9 | PA3872 | narI | respiratory nitrate reductase gamma chain | 3.21 | 0.01 | ↑ |
| 10 | PA2868 | NA | hypothetical protein | 3.16 | 0.01 | ↓ |
| 11 | PA3871 | NA | probable peptidyl-prolyl cis-trans isomerase, PpiC-type | 3.01 | 0.02 | ↑ |
| 12 | PA0532 | NA | hypothetical protein | 2.99 | 0.01 | ↓ |
| 13 | PA3395 | nosY | NosY protein | 2.98 | 0.02 | ↑ |
| 14 | PA0520 | nirQ | regulatory protein NirQ | 2.9 | 0.02 | ↑ |
| 15 | PA3873 | narJ | respiratory nitrate reductase delta chain | 2.89 | 0.04 | ↑ |
| 16 | PA0516 | nirF | heme d1 biosynthesis protein NirF | 2.84 | 0.02 | ↑ |
| 17 | PA2663 | ppyR | psl and pyoverdine operon regulator, PpyR | 2.82 | 0.05 | ↑ |
| 18 | PA0518 | nirM | cytochrome c-551 precursor | 2.82 | 0.03 | ↑ |
| 19 | PA0519 | nirS | nitrite reductase precursor | 2.74 | 0.03 | ↑ |
| 20 | PA0525 | NA | probable dinitrification protein NorD | 2.73 | 0.03 | ↑ |
| 21 | PA3392 | nosZ | nitrous-oxide reductase precursor | 2.72 | 0.03 | ↑ |
| 22 | PA4354 | NA | conserved hypothetical protein | 2.7 | 0.04 | ↓ |
| 23 | PA3393 | nosD | NosD protein | 2.6 | 0.05 | ↑ |
| 24 | PA3407 | hasAp | heme acquisition protein HasAp | 2.59 | 0.05 | ↑ |
| 25 | PA3874 | narH | respiratory nitrate reductase beta chain | 2.58 | 0.05 | ↑ |
| 26 | PA1942 | NA | hypothetical protein | 2.57 | 0.05 | ↓ |

Genes are arranged in decreasing order of Fold Change. Databases consulted for gene functions were: NCBI gene database (https://www.ncbi.nlm.nih.gov/nuccore/NC_002516); KEGG (Kyoto Encyclopaedia of Genes and Genomes: https://www.genome.jp/kegg/); Uniprot (https://www.uniprot.org/). NA: not available

### 2.13. Statistics

All results reported are means of three or more independent experiments, each performed in triplicate. Statistical significance was assessed through t-test performed using Microsoft Excel^®^, and data with p≤0.05 was considered to be significant.

## 3. Results and Discussion

### 3.1. Silver inhibits growth of *P. aeruginosa* and modulates QS-regulated pigment formation

*P. aeruginosa* was challenged with different Silversol^®^ concentrations (0.03- 2 ppm). Bacterial growth and pyoverdine production remained unaffected till 0.5 ppm Silversol^®^, while pyocyanin formation was inhibited in a dose-dependent fashion 0.5 ppm onward (Figure 1). Effect of Silversol^®^ on pyoverdine production appeared to follow an inverted U-shaped hormetic dose response curve (Calabrese, 2014) between the concentration range 0.5-1.5 ppm. Since Silversol^®^ inhibited pyocyanin formation at 0.5 ppm without affecting bacterial growth, this concentration can be said to be purely quorum modulatory. Since 1 ppm seemed to be the ∼IC_50_, and this concentration of Silversol^®^ also had a heavy effect on both quorum-regulated pigments, we chose this as the test concentration for further experiments. While Silversol^®^ seemed to inhibit pigment production completely 1.5 ppm onward, and caused complete visible inhibition of growth 2 ppm onward, the MBC was found to be somewhere between 16-20 ppm (Figure S3). When cells from experimental tubes were plated onto Pseudomonas agar (not containing any Silversol**^®^**), only those coming from 1 ppm silver tubes gave pigmented growth, while those from 10-15 ppm silver tubes grew without pigmentation. Inoculum coming from the 20 ppm silver tubes failed to give rise to any growth on agar plates indicating that MBC has been reached.

**Figure 1.**
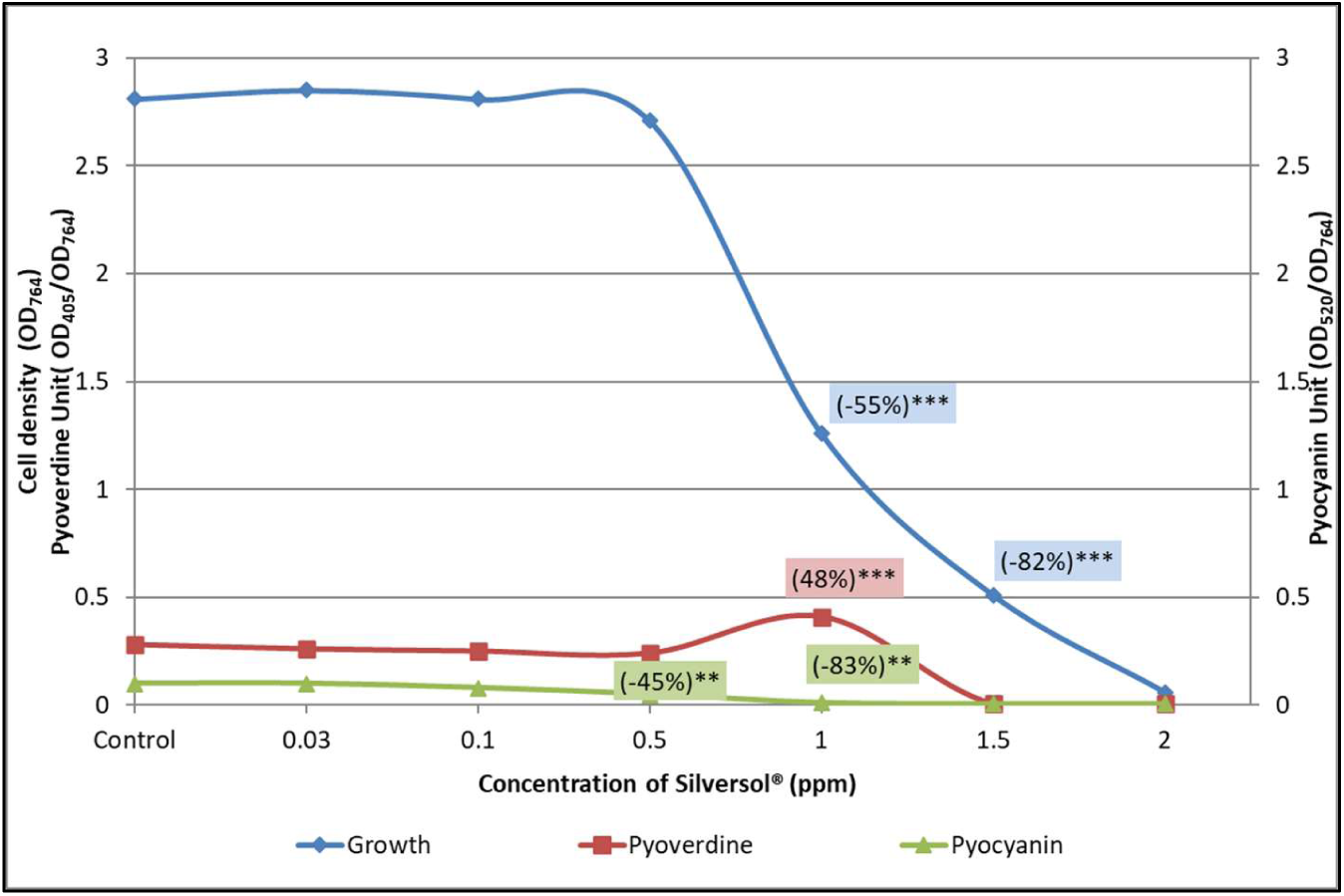
Growth-inhibitory and quorum-modulatory effect of Silversol^®^ on *P. aeruginosa.* Bacterial growth was measured as OD764; OD of pyoverdine and pyocyanin was measured at 405 nm and at 520 nm respectively. Pyoverdine Unit and Pyocyanin Unit were calculated as the ratio OD405/OD764 and OD520/OD764(an indication of pigment production per unit of growth); Ofloxacin (0.3 μg/mL) inhibited growth by 76**%±0.1 with affecting pigment production. **p<0.01, ***p<0.001; minus sign (-) in parentheses indicate a decrease over control.

### 3.2. Silver appears to disturb iron homeostasis of *P. aeruginosa*

Silversol^®^ (1 ppm) enhanced pyoverdine production by almost 1.5-fold, and since pyoverdine is a siderophore (Kang et al., 2018), we speculated that Silversol^®^ might be disturbing iron-homeostasis in *P. aeruginosa*. To have additional insight into this aspect, we inoculated blood agar plates with silver-exposed *P. aeruginosa*, and these cells were found to be more haemolytic than their silver-not-exposed counterparts (Figure 2). Taking the effect of silver on pyoverdine production and haemolytic activity of the pathogen together, it can be said that silver-exposed cells seem to experience iron-limitation and to overcome this the cells are producing more siderophore and haemolysis.

**Figure 2:**
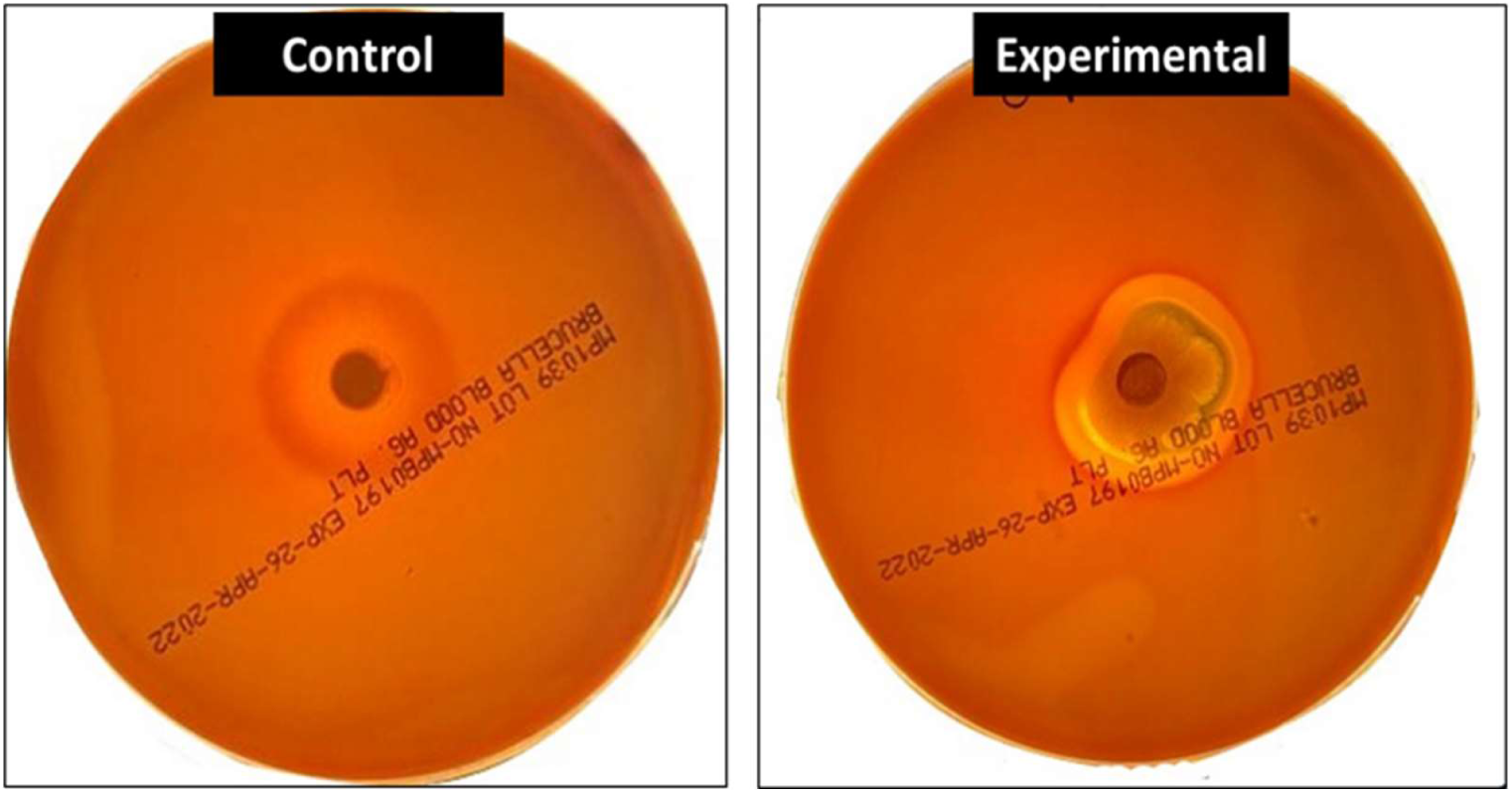
Silver pre-treated *P. aeruginosa* exhibits enhanced haemolysis on blood agar plates. Silver-pretreated *P. aeruginosa* when inoculated onto blood agar plates, produced a clearer zone of haemolysis than its control counterpart receiving no previous exposure to silver.

### 3.3. Silver-treated *P. aeruginosa* displays higher susceptibility to augmentin and streptomycin

To investigate whether silver exposure influences antibiotic susceptibility of 1. *P. aeruginosa*, we grew *P. aeruginosa* in presence of 1 ppm Silversol^®^, and the antibiogram of the resulting cells (after washing with phosphate buffer) was compared to that of control cells. Silver-pre-exposed cells displayed a phenotype change from ‘resistant’ to ‘sensitive’ against augmentin; and a notable increase (38%) in susceptibility to streptomycin (Table-2; Figure 3). Such resistance-modifying activity of the silver becomes quite important given *P. aeruginosa* is known rarely to be susceptible to augmentin (Deforges et al., 1985), and augmentin resistance is quite prevalent among its clinical isolates (Khan et al., 2022). Streptomycin-resistance was reported by Cervantes-Vega et al (1986) to be among the most common resistance patterns in *P. aeruginosa* clinical isolated from Mexico.

**Figure 3:**
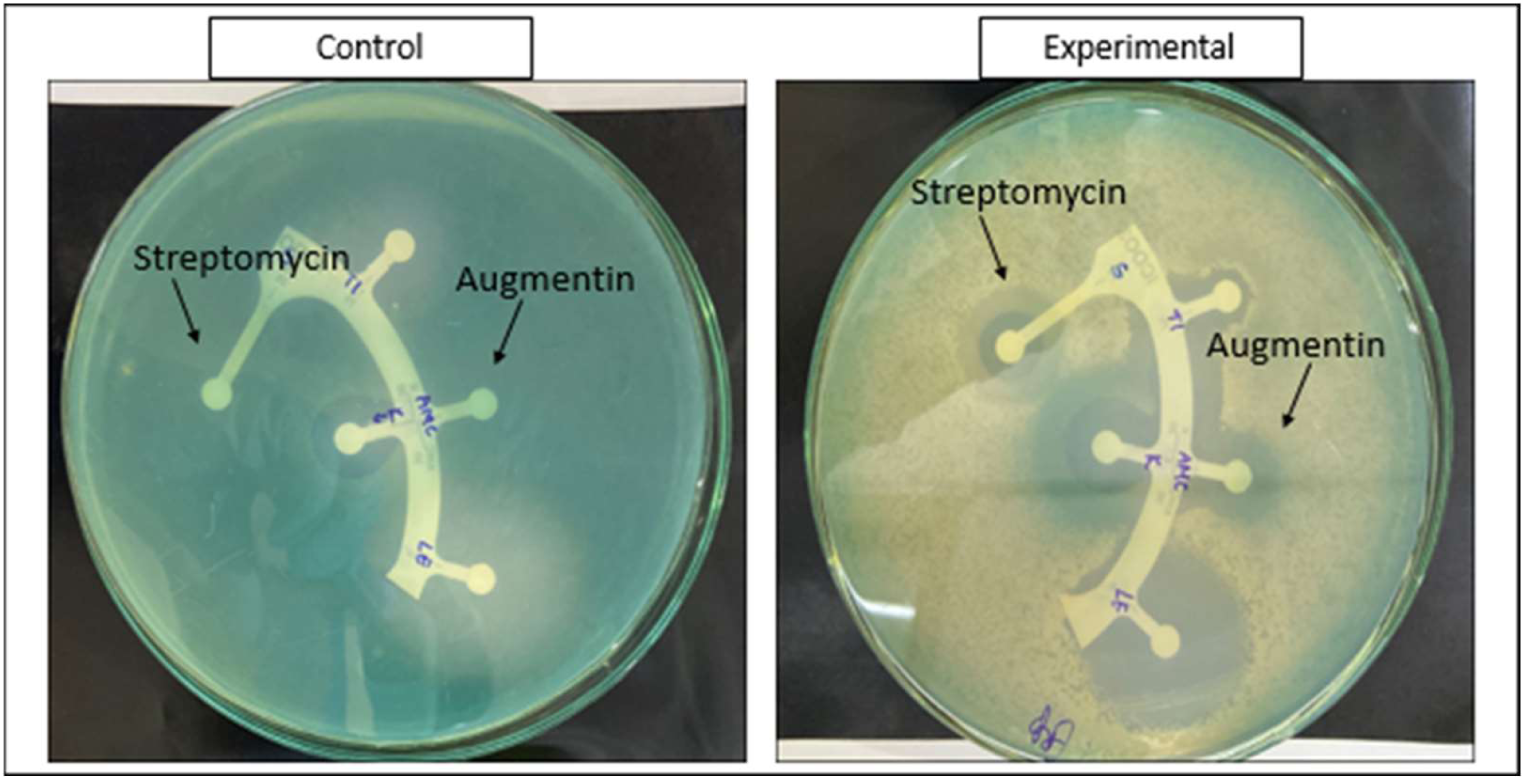
Silver pre-treated enhances *P. aeruginosa*’s susceptibility to augmentin and streptomycin. Silver-pretreated cells seem to be altered in their response to antibiotics with respect to growth as well as QS-regulated pigmentation. On these plates of cation-adjusted Muller-Hinton agar, clear zones of inhibition can be seen surrounding the discs of augmentin and streptomycin in the experimental plate inoculated with silver-pre-treated cells, while such zones are absent (augmentin) or faint (streptomycin) from the control plate inoculated with *P. aeruginosa* having no previous silver-exposure.

### 3.4. Silversol^®^ could partially eradicate pre-formed *P. aeruginosa* biofilm

While Silversol^®^ inhibited biofilm formation in *P. aeruginosa* partially (19%), it could also eradicate the pre-formed biofilm (by 34%) and reduce biofilm viability by 30% (Figure 4). Since bacterial biofilms are known for displaying higher antibiotic resistance than their planktonic counterparts (Gupta et al., 2018), biofilm-eradicating agents can help restoring the antibiotic susceptibility of bacterial cells by freeing them of the biofilm matrix.

**Figure 4.**
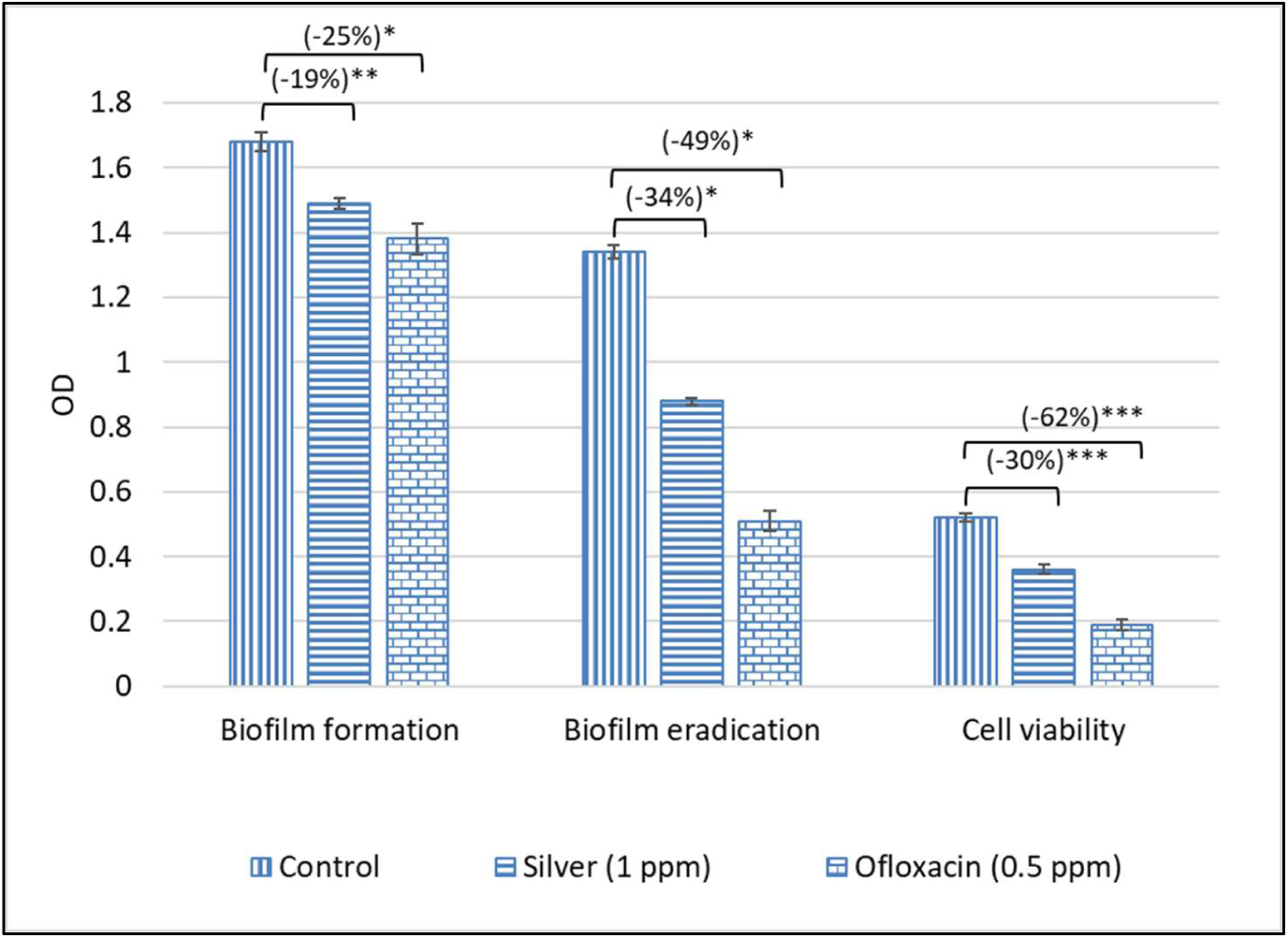
Silver could partially inhibit *P. aeruginosa* biofilm formation, and eradicate the pre-formed biofilm. Crystal violet assay was performed to quantify biofilm formation, and eradication. Cell viability in the biofilm was estimated through MTT assay. *p<0.05, **p<0.01, ***p<0.001; minus sign (-) in parentheses indicate a decrease over control.

Since exopolysaccharides (EPS) constitute an important component of bacterial biofilm matrix, we quantified EPS in control vs. silver-treated *P. aerugin*osa culture. Though the absolute EPS in experimental and control cultures was at par, nullifying it against reduced cell density in silver-exposed culture, EPS production in experimental culture was found to be enhanced by 76.34% (Figure 5). Higher EPS synthesis can be considered as an indication of envelope stress (Mohammed et al., 2020) in the silver-exposed bacteria. When faced with a variety of stress conditions, *P. aeruginosa* is known to respond by escalating its EPS synthesis as a survival strategy, since EPS serves as a protective matrix (O’Toole et al., 2000; Yang et al., 2008), whose production is upregulated during nutrient limitation and oxidative stress (Hu et al., 2019; Morales et al., 2012). This study found the sub-MIC level of Silversol to disturb QS-regulated traits in *P. aeruginosa* such as pigmentation (Figure 1) and biofilm formation (Figure 4), and the quorum sensing system in *P. aeruginosa* is known to influence EPS production too (Davies et al., 1998). Studying the correlation between EPS production and stress-response in *P. aeruginosa* can provide new insights into its survival mechanisms and pathogenicity.

**Figure 5.**
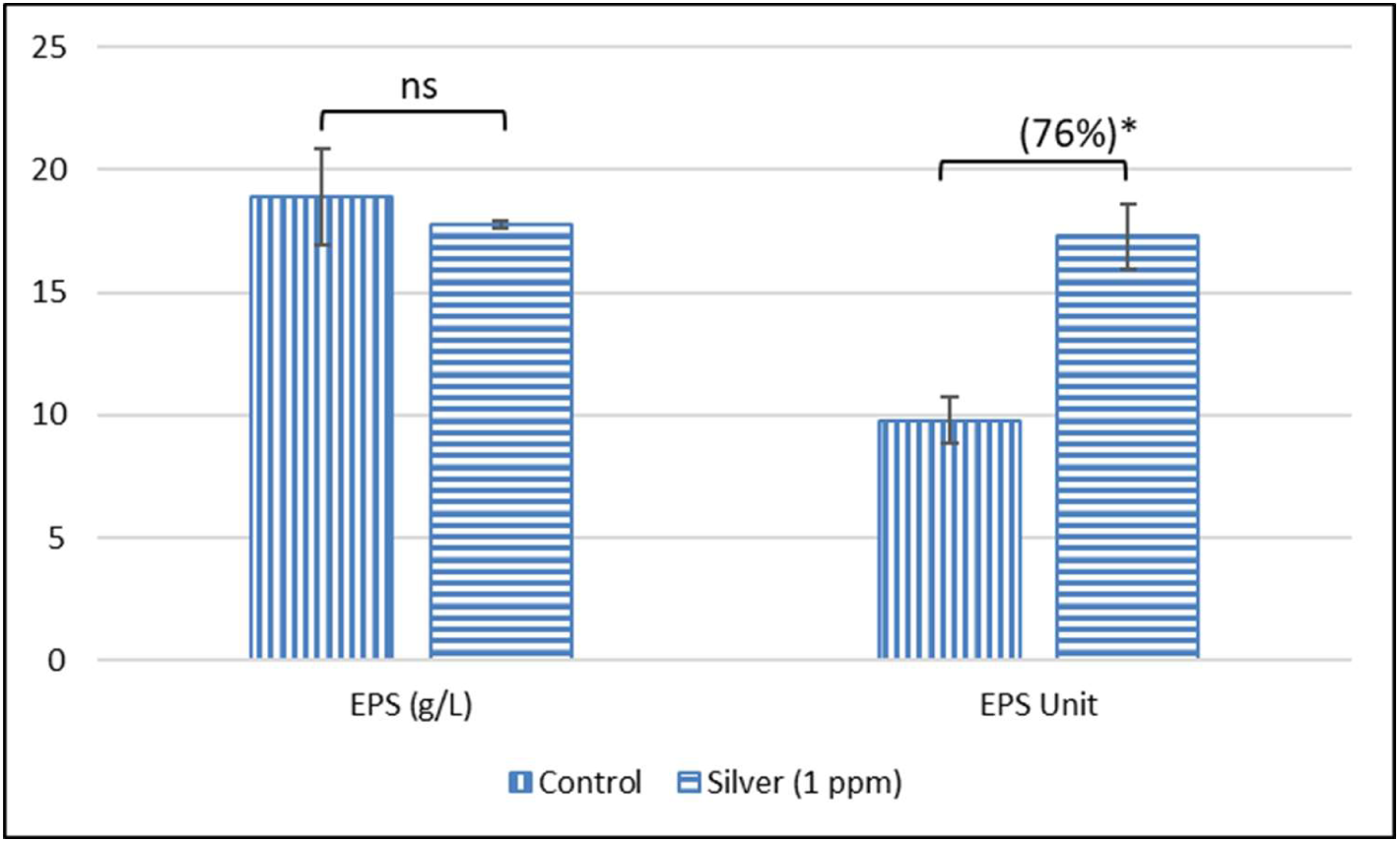
Silver enhances EPS synthesis in *P. aeruginosa*. Though there are lesser number of cells in the silver-supplemented media, they synthesized EPS in amount equal to their silver-unexposed counterparts. EPS Unit was calculated as Cell Density (OD764): EPS (g/L) ratio. *p<0.05; ns: non-significant

### 3.5. *P. aeruginosa* seemed to upregulate its protein synthesis in presence of silver

*P. aeruginosa* exposed to sub-MIC of Silversol^®^ was found to have higher intracellular as well as extracellular protein concentration than its silver-non-exposed counterpart (Figure 6). Bacterial response to the known inhibitor of protein synthesis (Kanamycin) was also similar. It may be said that bacteria responds to the inhibitory effect of such antimicrobials exerting their action through suppressing protein synthesis, by upregulating its protein sysnthesis and/or secretion machinery to compensate the inhibitory effect of the antimicrobial agents (Singh et al., 2001). Such upregulation of protein synthesis might have stemmed from the translational reprogramming in the stressed cells (Liu and Qian, 2014), as they are facing the stress of the antibacterial activity of the silver.

**Figure 6.**
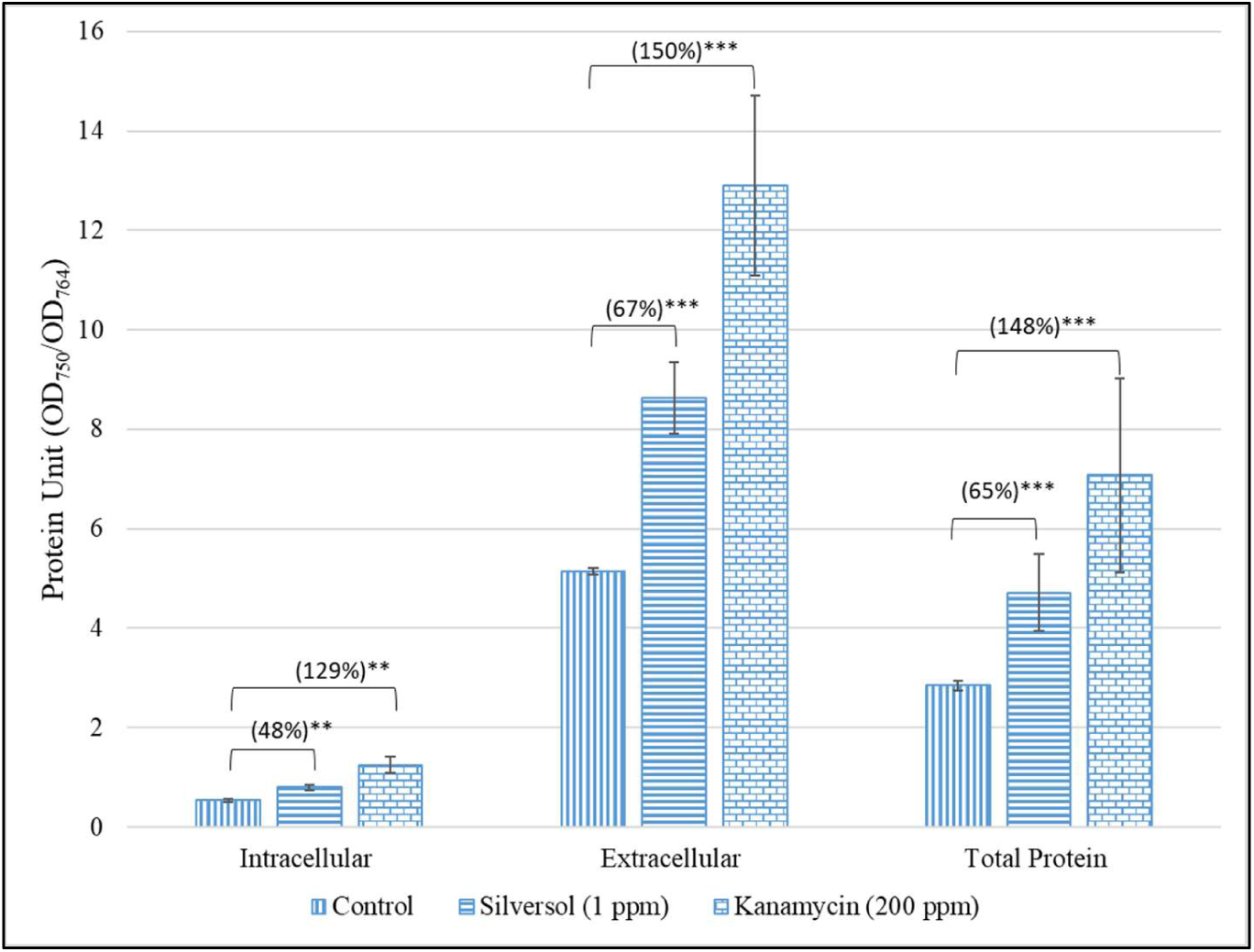
*P. aeruginosa* grown in presence of Silversol^®^ registered higher protein synthesis. Intracellular and extracellular protein concentrations in *P. aeruginosa* grown in presence of Silversol^®^ at sub-MIC level were significantly higher as compared to its silver-non-exposed counterpart. Kanamycin employed as a positive control at its sub-MIC level also generated similar response from bacterial culture. Protein Unit (i.e., Protein concentration: Cell density ratio) was calculated to nullify any effect of cell density on protein production. **p<0.01, ***p<0.001

### 3.6. Silver induces nitrosative stress in *P. aeruginosa*

Since denitrification pathway is an important metabolic pathway in *P. aeruginosa*, and enzymes involved in detoxification of reactive nitrogen species are proposed to be potential anti-pathogenic targets (Joshi et al., 2019; Ruparel et al., 2023), we quantified one of the intermediates of denitrification pathway, nitrite (NO_2_^-^), in silver-exposed *P. aeruginosa*. Culture supernatant of the latter was found to possess 37% higher nitrite (Figure 7) than that grown in absence of silver. Higher nitrite concentrations can have multiple effects on *P. aeruginosa* physiology and virulence. It can exert its toxicity by disrupting electron transport chain, and can also impair bacterial virulence, besides modulating susceptibility to various antibiotics. Nitrite can react with other reactive nitrogen species (RNS) to form highly reactive intermediates, such as peroxynitrite (ONOO^-^), which can damage cellular components, including proteins, lipids, and DNA (Poole, 2005). Nitrite can interfere with the electron transport chain in *P. aeruginosa*, leading to a decrease in ATP production and compromised energy metabolism (Heales et al., 1999; Wesselink et al., 2019). This disruption can negatively impact various cellular processes and growth. Nitrite can also inhibit the production of virulence factors such as pyocyanin, elastase, and siderophores (Schreiber et al., 2006). Nitrite build-up can negtively impact bacterial physiology by trigerring the overexpression of efflux pumps and altering membrane permeability (Ciofu, 2003).

**Figure 7.**
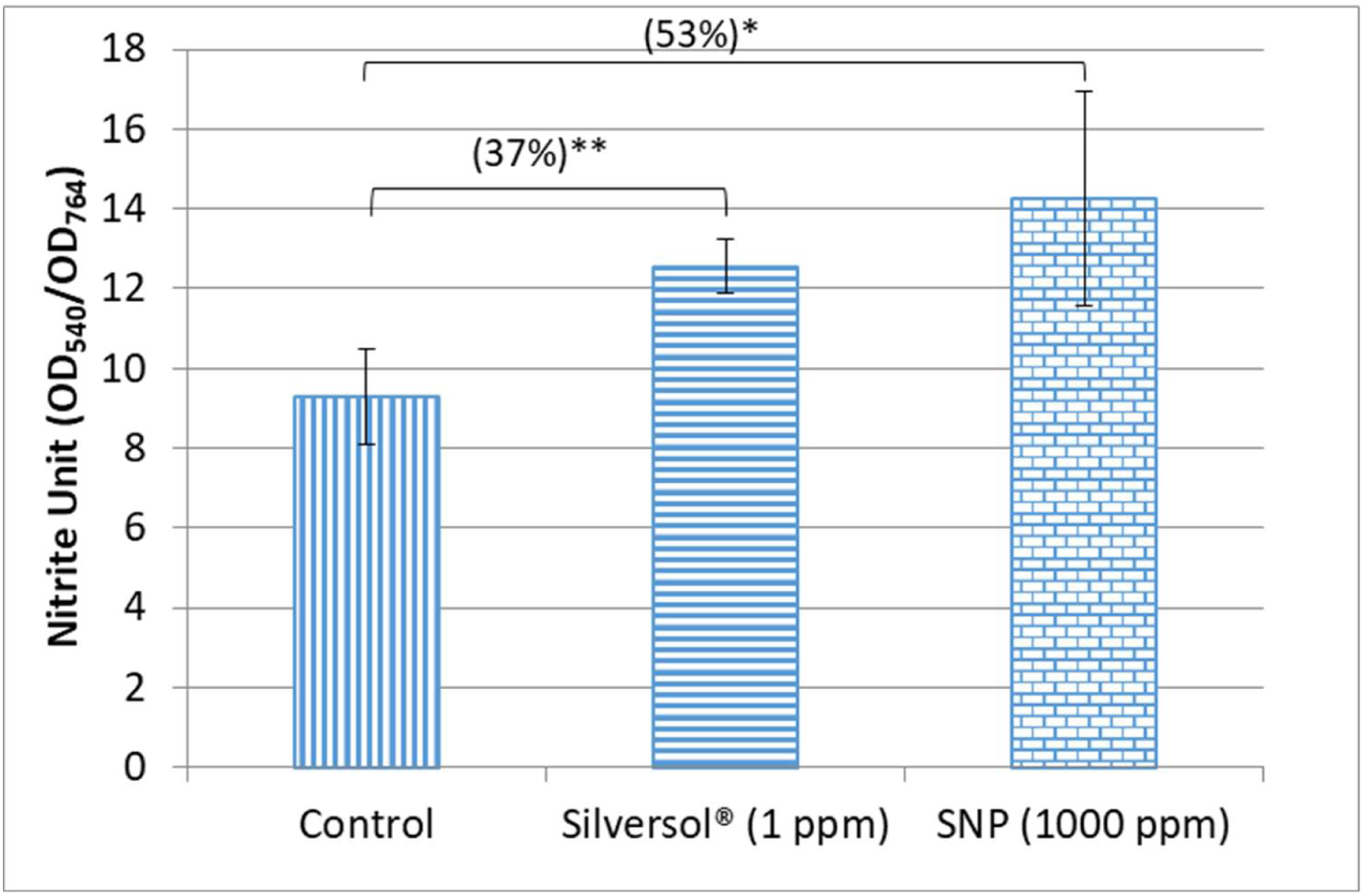
Silversol^®^-treated *P. aeruginosa* culture has higher extracellular accumulation of nitrite. Silversol^®^ caused nitrite concentration in *P. aeruginosa* culture supernatant to rise when compared to control. Sodium nitroprusside (SNP) used as a positive control also caused higher nitrite build up in *P. aeruginosa* culture. Nitrite Unit (i.e., Nitrite concentration: Cell density ratio) was calculated to nullify any effect of cell density on nitrite production. *p<0.05, **p<0.01

**Figure 8.**
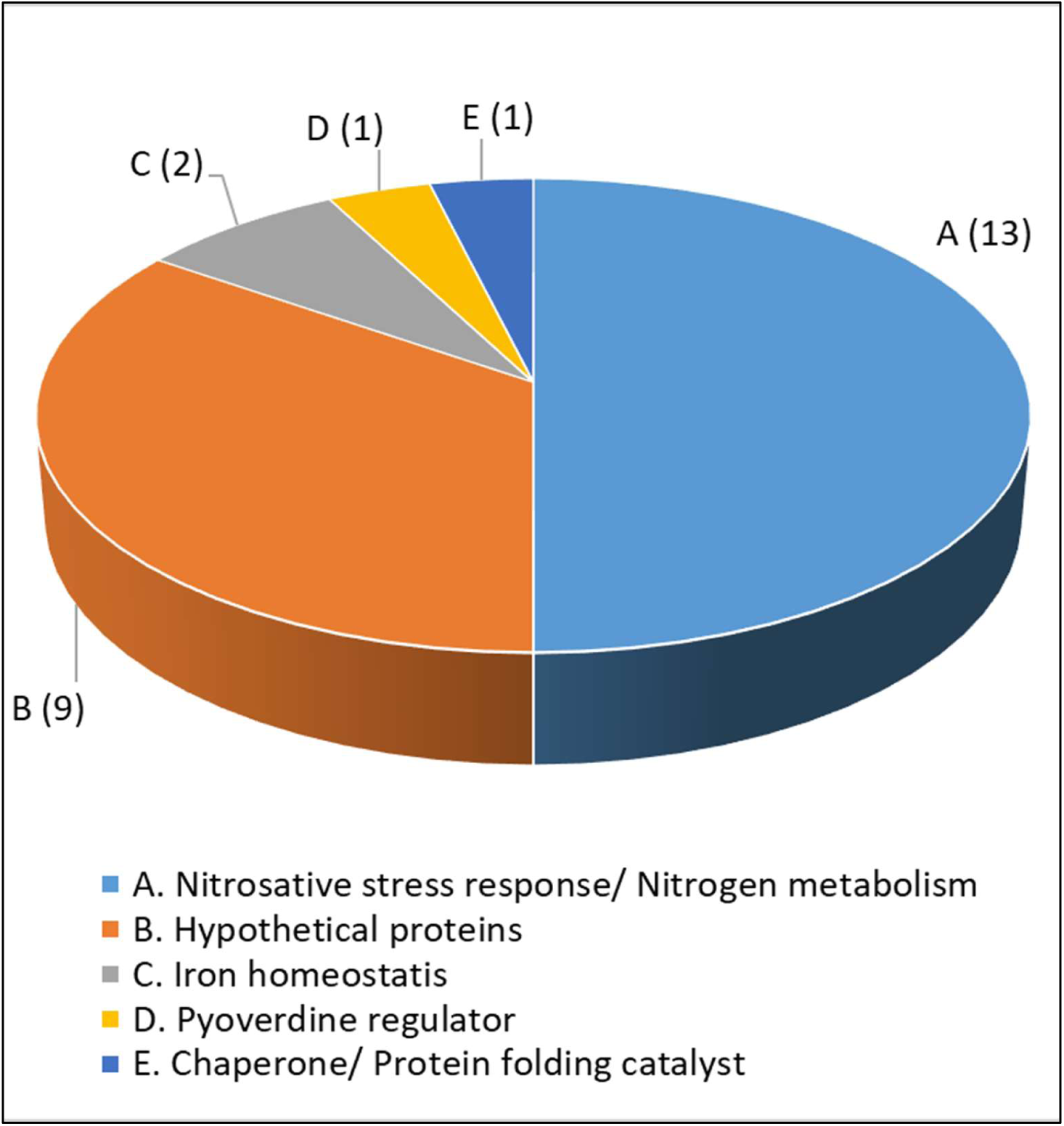
Function-wise categorization of the significantly differentially expressed genes in Silver-treated *Pseudomonas aeruginosa.* Figures in parentheses indicate number of genes in that particular category.

### 3.7. Differential gene expression in silver-treated *P. aeruginosa*

Silversol^®^-exposed *P. aeruginosa* when subjected to whole transcriptome analysis, it was found to express a total of 26 genes (0.48% of total genome) differently (log fold-change ≥ 2 and FDR ≤ 0.05). List of differentially expressed genes (DEG) is provided in Table-3, and corresponding heat map (Figure S4) and volcano plot (Figure S5) can be seen in supplementary file. When we looked for functions of the DEG in appropriate databases like KEGG, PDB, or UniProt, all the 6 downregulated genes turned out to be hypothetical proteins. Amongst the upregulated genes in silver-exposed *P. aeruginosa* culture, 13 genes were associated with nitrogen metabolism, 2 were associated with heme biosynthesis or acquisition, three were hypothetical proteins, 1 was pyoverdine regulator and 1 was a protein folding catalyst. Function-wise categorisation of all DEG is presented in Figure-8.

Empirically the transcriptomic profile indicated Silversol^®^ to disturb iron homeostasis and nitrogen metabolism in *P. aeruginosa*. The hypothesis of disturbance of iron homeostasis corroborates well with results of our *in vitro* experiments for assessing haemolytic potential (Figure 2) and pyoverdine (Figure 1) production. It seems that in presence of Silversol^®^, the bacteria are facing iron-limitation and is forced to upregulate the genes coding for heme synthesis and siderophore production to maintain iron homeostasis. This observation is important in light of the fact that iron is an essential micronutrient for bacteria, and since the concentration of free iron in human serum is quite low (Litwin and Calderwood, 1993), pathogens have to develop iron-scavenging strategies for a successful in-host survival. Differential expression of multiple genes associated with nitrogen metabolism also corroborates well with increased nitrite concentration in silver-treated bacterial culture supernatant (Figure 7).

To have a holistic idea of the antibacterial mechanism of Silversol^®^, we subjected all the DEG to network analysis through STRING. The resulting protein-protein interaction (PPI) (Figure 9) network had 26 nodes connected through 74 edges with an average node degree of 5.69. Since the number of edges (74) in this PPI network is almost 25-fold higher than expected (03) with a PPI enrichment *p*–value <1.0e^-^16, this network can be said to possess significantly more interactions among the member proteins than what can be expected for a random set of proteins having identical sample size and degree distribution. Such an enrichment is suggestive of the member proteins being at least partially biologically connected.

**Figure 9.**
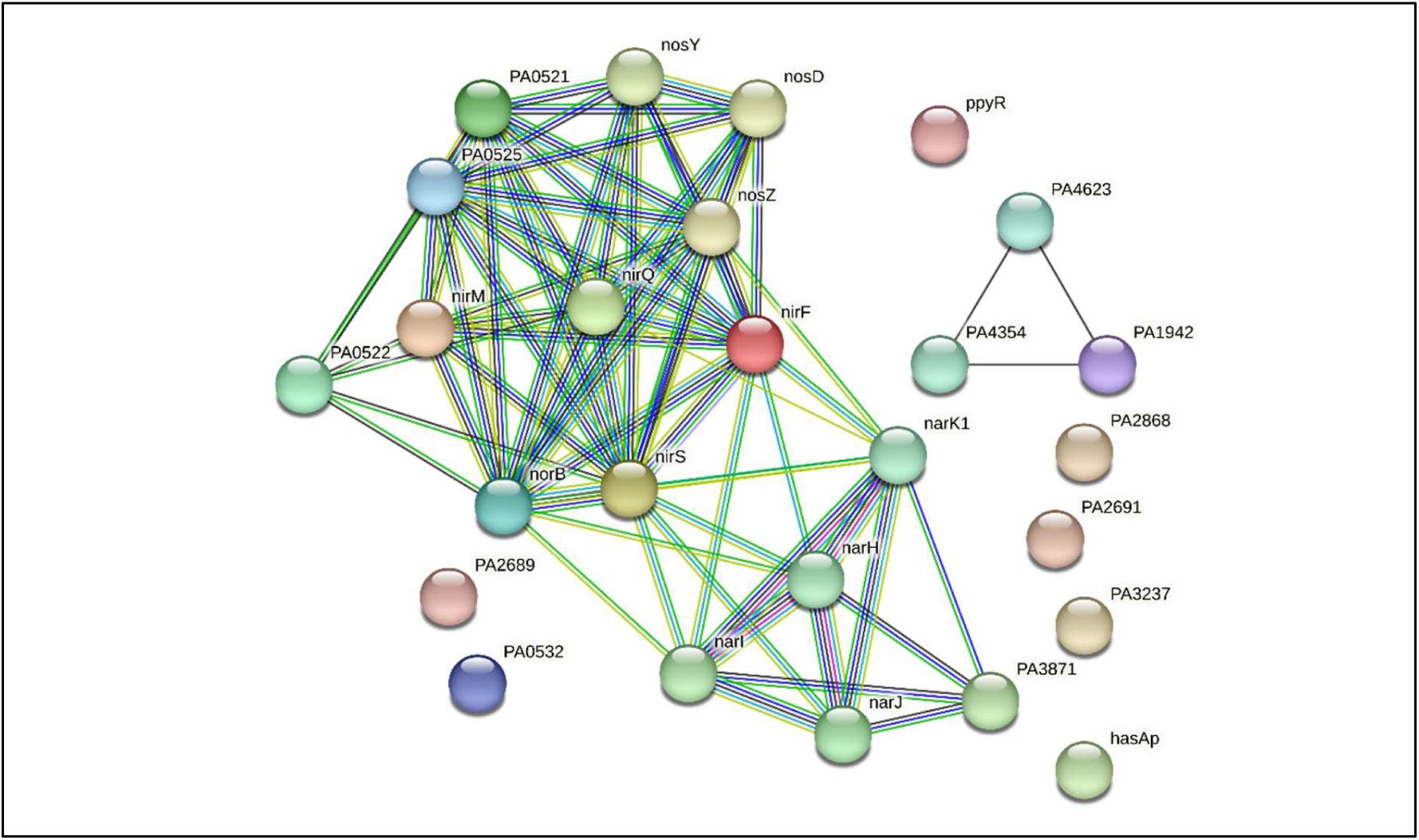
PPI network of DEGs in Silversol^®^-exposed *P. aeruginosa*.

When we arranged all the 26 nodes in decreasing order of node degree, 19 nodes were found to have a non-zero score, and we selected top 16 genes with a node degree ≥4 (Table 4) for further ranking by different cytoHubba methods. Then we looked for genes which appeared among top ranked candidates by ≥6 cytoHubba methods. This allowed us to shortlist ten genes (Table 5) which were ranked among top-10 by ≥9 cytoHubba methods to be taken for further cluster analysis. Interaction map of these 10 important genes (Figure 10) showed them to be networked with the average node degree score of 8. Number of edges possessed by this network was 40 as against expected 0 for any such random set of proteins. These 10 genes were found to be distributed among five different local network clusters. Strength score for each of these clusters was >1.86. While five of the proteins (norB, PA0525, nirS, nirQ, and nosZ) were members of all the five clusters, two proteins (nosY and PA0521) were part of four different clusters. Of the remaining three, nirF was part of three different clusters, while nirM and narK1 were respectively members of two and a single cluster. The multi-cluster proteins identified here can be said as not only the most important targets of Silversol^®^, but also hub proteins with high network centrality and potential targets for novel anti-Pseudomonas drug discovery programmes.

**Figure 10.**
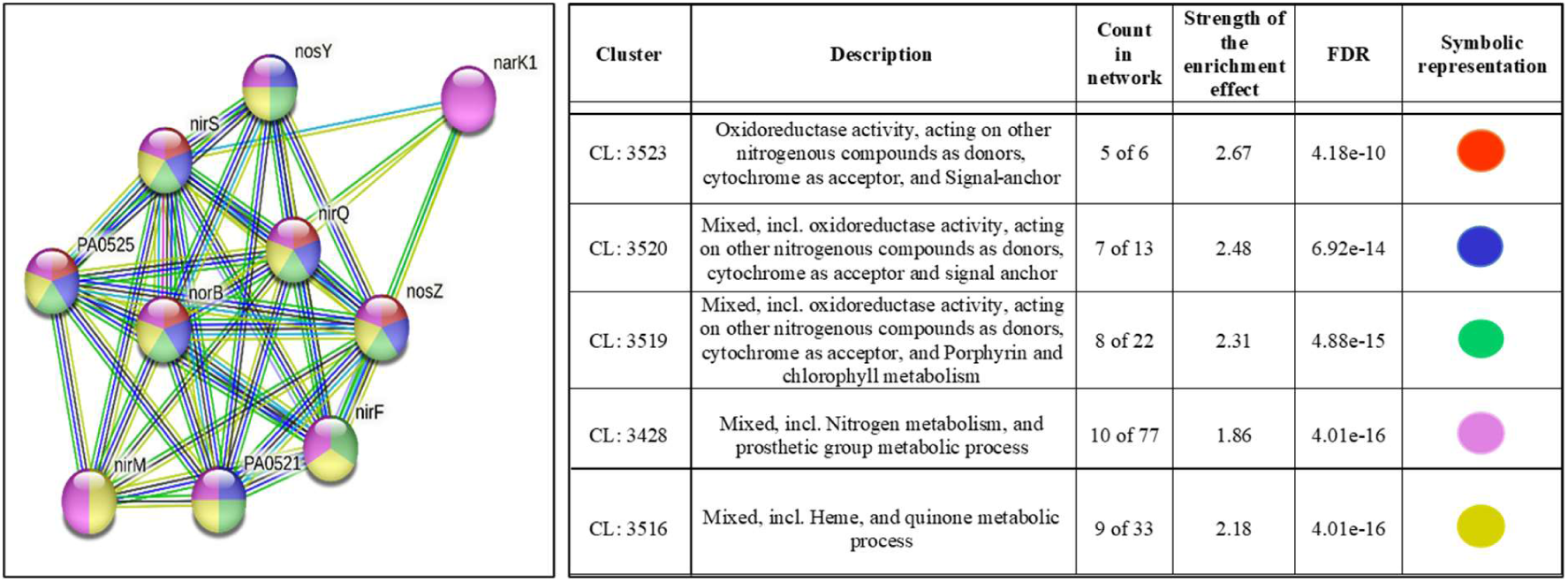
PPI network of top-ranked genes revealed through cytoHubba among DEG in Silversol^®^*-*exposed *P. aeruginosa*.

**Table 4.** Node degree score of the genes mentioned in Table 3.

| Sr. No. | Gene ID/Symbol | Node Degree |
| --- | --- | --- |
| 1 | nirS | 14 |
| 2 | norB | 13 |
| 3 | nirF | 12 |
| 4 | nirQ | 11 |
| 5 | PA0521 | 10 |
| 6 | norD | 10 |
| 7 | nosZ | 10 |
| 8 | narK1 | 9 |
| 9 | nirM | 8 |
| 10 | nosD | 8 |
| 11 | nosY | 8 |
| 12 | narH | 7 |
| 13 | narI | 7 |
| 14 | PA0522 | 6 |
| 15 | narJ | 5 |
| 16 | PA3871 | 4 |
| 17 | PA1942 | 2 |
| 18 | PA4354 | 2 |
| 19 | PA4623 | 2 |

**Table 5.**
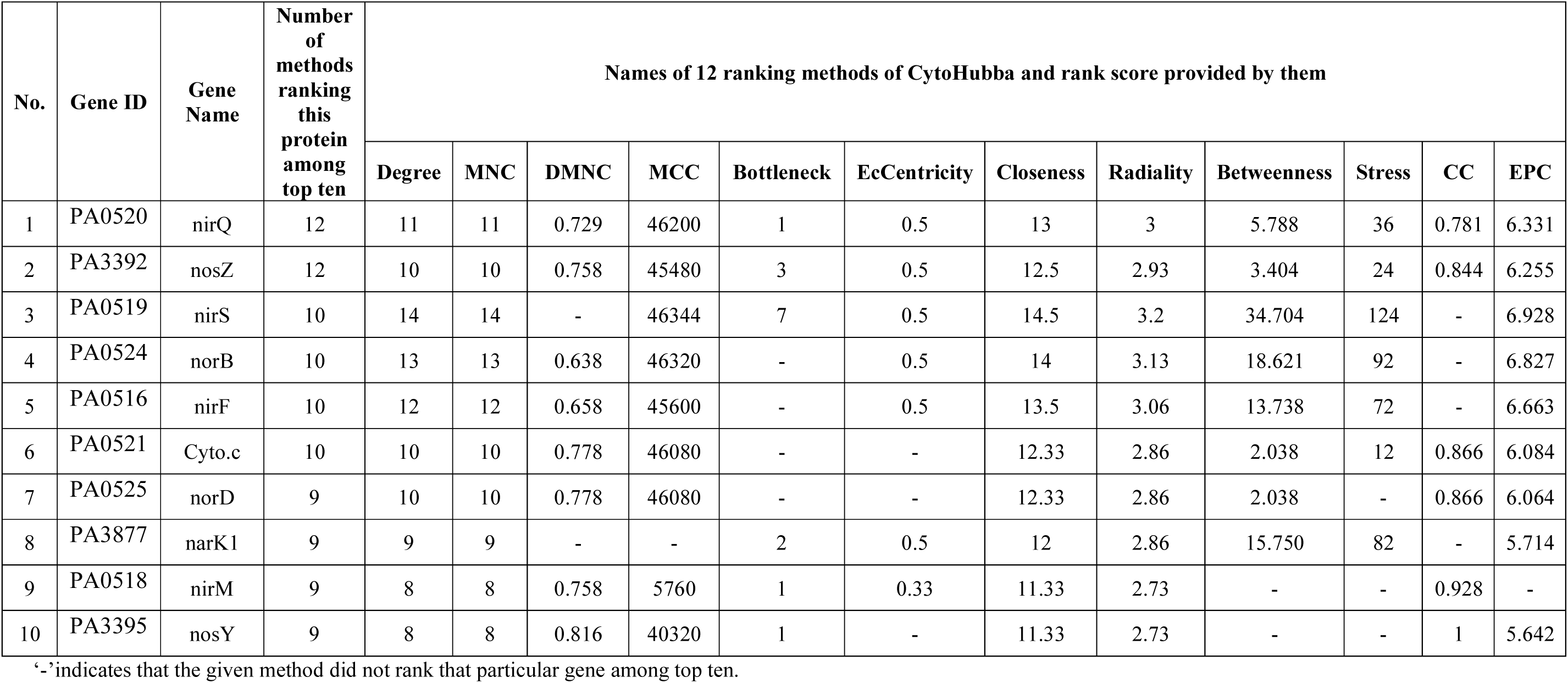
Top ten cytoHubba-ranked genes from among the top-16 with respect to node degree.

We also conducted a gene cooccurrence pattern analysis of gene families across genomes (through STRING) with respect to the potential hubs identified by us (Figure 11). None of the 10 potential targets was shown to be present in humans, hence any anti-pathogenic drugs capable of causing dysregulation of these genes are least likely to toxic to humans. This corroborates well with reports showing Silversol^®^ to be safe for human consumption (de Souza et al., 2021). Three of the Silversol^®^’s target genes in *P. aeruginosa* were also shown to be present in *Staphylococcus aureus*. However, except narK1, no other target proteins were shown to be present in other important pathogens. Since Silversol^®^ is already known to be active against a wide spectrum of gram-positive and gram-negative bacteria (de Souza et al., 2006), it may be believed to be acting against different organisms through different mechanisms.

**Figure 11.**
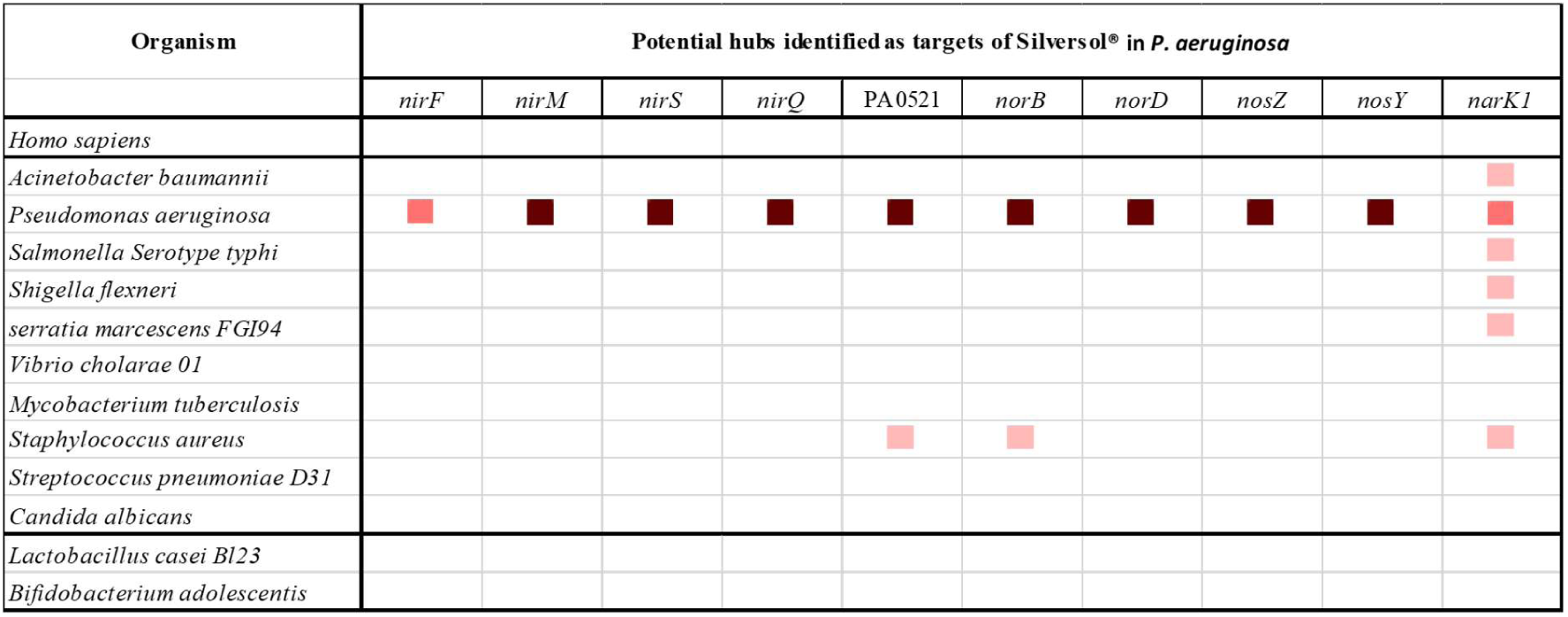
Cooccurrence analysis of genes coding for potential targets in *P. aeruginosa*.

One of the side-effect with conventional antibiotics is that they fail to differentiate between the ‘good’ (symbionts in human microbiome) and ‘bad’ (pathogens) bacteria, and hence their therapeutic use may lead to gut dysbiosis. An ideal antimicrobial agent is expected to target pathogens exclusively without causing gut dysbiosis. In this respect, a target in pathogenic bacteria absent from symbionts of human microbiome will be most suitable candidate for antibiotic discovery programmes. To have some insight on this front with respect to the targets identified by us, we run a gene cooccurrence analysis with two representative ‘good’ bacteria too, reported to be part of healthy human microbiome, and the said targets were not shown to be present in them. This corroborates with the selective inaction of Silversol on probiotic strains of *Lactobacillus acidophilus* and *Bifidobacterium longum* documented in internal reports of Viridis Biopharma (Selective inaction of ASAP on probiotics. Unpublished raw data, 2004).

### 3.8. Target validation through RT-PCR

Based on the network analysis of transcriptome of silver-treated *P. aeruginosa*, we selected following three genes for further validation through RT-PCR: nor B (identified as member of all the 5 clusters in network analysis), PA0521 (identified as member of 4 different clusters in network analysis), and narK1 (shown by gene cooccurrence analysis to be present across multiple pathogenic genera); wherein they were found to be up-regulated in silver-treated *P. aeruginosa* by 2.65-fold, 2.32-fold, and 7.37-fold respectively (Figure 12).

**Figure 12.**
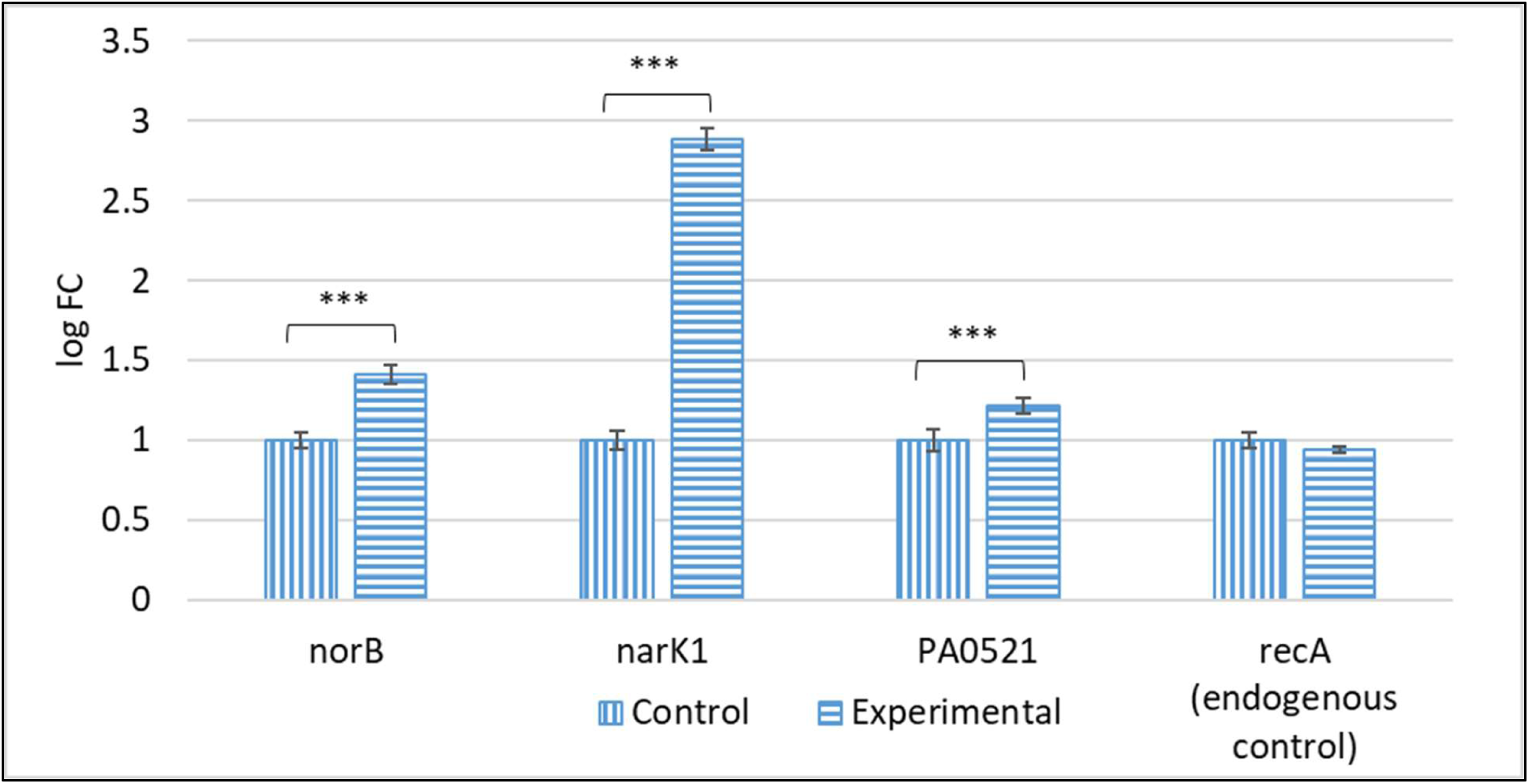
Confirmation of differential expression of selected genes in Silversol-treated *P. aeruginosa* through RT-PCR. ***p<0.001

## 4. Conclusion

Silver has long been known for its antimicrobial activity. Though literature contains reports describing a variety of modes of antibacterial action of silver and its nanoparticles, our knowledge on this front is far from complete. Antibacterial mechanisms of silver previously described by various researchers include generation of oxidative stress (Lee et al., 2014; Ahn et al., 2014; Hande et al., 2021), generating nitrosative stress (Hande et al., 2021), disruption of membrane integrity (Morones-Ramirez et al., 2013), protein denaturation (McQuillan and Shaw, 2014; Yin et al., 2020), inhibition of enzymatic activity (Li et al., 2010; Sinlo et al., 2014), interference with quorum sensing (Subhadra et al., 2018; Qais et al., 2020), inducing DNA damage (Ahn et al., 2014) and interfering with its replication and transcription (Feng et al., 2000), inhibition of efflux pumps (Subhadra et al., 2018), etc. The present study investigated effect of sub-MIC level of colloidal silver on *P. aeruginosa*’s growth, QS-regulated pigmentation, biofilm, protein synthesis, nitrogen metabolism, EPS synthesis, haemolytic activity and siderophore production, antibiotic susceptibility, and gene expression at the whole transcriptome level. A schematic summary of Silversol’s multiple effects on *P. aeruginosa* is presented in Figure 13. Disruption of iron homeostasis and generation of nitrosative stress seemed to be the major mechanisms of anti-*P. aeruginosa* activity of silver in this study. Differential expression of three important genes involved in denitrification pathway in silver-exposed *P. aeruginosa* was confirmed through RT-PCR too. Hub proteins identified in this study as major targets of silver in *P. aeruginosa* warrants further investigation with respect to validating their targetability e.g., by confirming defective growth of mutant strains of *P. aeruginosa* bearing deletion of one or more of the identified hub genes. Such validated targets can prove vital to various antibiotic discovery efforts globally. Though the gene expression profile of the bacterium under influence of sub-lethal concentrations of silver may vary from that under influence of its lethal concentrations, this study provides useful insights into antibacterial mechanism of silver and identification of its potential targets in *P. aeruginosa*.

**Figure 13.**
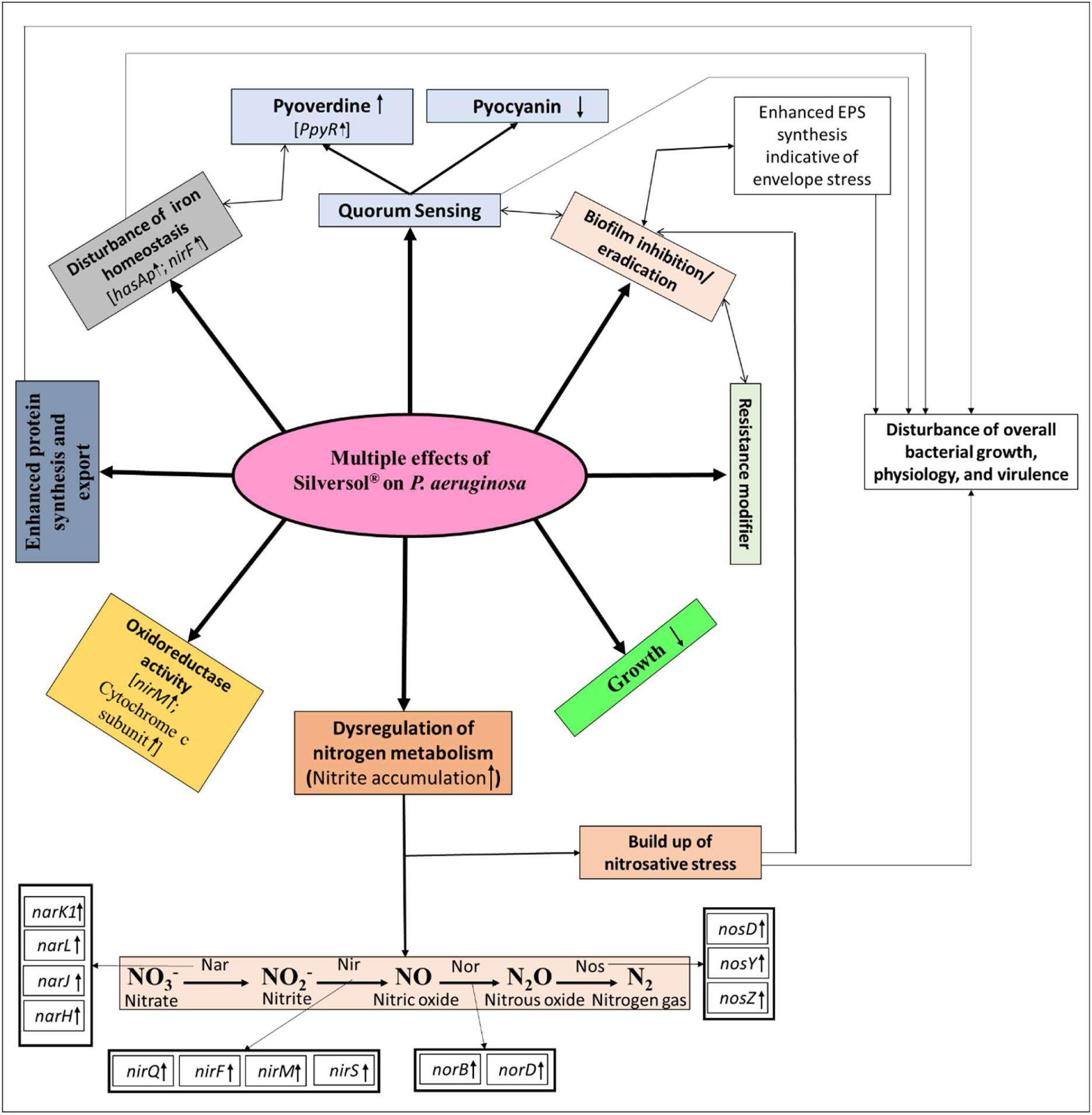
Overall schematic of multiple effects of Silversol^®^ on *Pseudomonas aeruginosa*. Various physiological and virulence traits of *P. aeruginosa* affected under the influence of sub-lethal concentration of Silversol**^®^** are depicted. Up (↑) or down (↓) regulation of the genes relevant to those traits is also indicated. EPS: exopolysaccharide

## Acknowledgement

Authors thank Nirma Education and Research Foundation (NERF), Ahmedabad for infrastructural support.

## References

1. Ahmad A, Wei Y, Syed F, Tahir K, Rehman AU, Khan A, Ullah S, Yuan Q. The effects of bacteria-nanoparticles interface on the antibacterial activity of green synthesized silver nanoparticles. Microb. Pathog. 2017;102:133–42. https://doi.org/10.1016/j.micpath.2016.11.030

2. Ahn JM, Eom HJ, Yang X, Meyer JN, Choi J. Comparative toxicity of silver nanoparticles on oxidative stress and DNA damage in the nematode, Caenorhabditis elegans. Chemosphere. 2014; 108:343–52. https://doi.org/10.1016/j.chemosphere.2014.01.078

3. Anandaradje A, Meyappan V, Kumar I, Sakthivel N. Microbial synthesis of silver nanoparticles and their biological potential. Nanopart. Med. 2020:99–133. DOI: 10.1007/978-981-13-8954-2_4

4. Andrews, S. FastQC: a quality control tool for high throughput sequence data. 2010.

5. Auger C, Lemire J, Cecchini D, Bignucolo A, Appanna VD. The metabolic reprogramming evoked by nitrosative stress triggers the anaerobic utilization of citrate in Pseudomonas fluorescens. PLoS One. 2011;6(12):e28469. https://doi.org/10.1371/journal.pone.0028469

6. Balachandar R, Navaneethan R, Biruntha M, Kumar KK, Govarthanan M, Karmegam N. Antibacterial activity of silver nanoparticles phytosynthesized from Glochidion candolleanum leaves. Mater. Lett. 2022; 311:131572. https://doi.org/10.1016/j.matlet.2021.131572

7. Calabrese EJ. Hormesis: a fundamental concept in biology. Microb. Cell. 2014;1(5):145. doi:10.15698/mic2014.05.145

8. Cervantes-Vega C, Chávez J, Rodríguez MG. Antibiotic susceptibility of clinical isolates of Pseudomonas aeruginosa. Antonie Van Leeuwenhoek. 1986;52:319–24. https://doi.org/10.1007/BF00428643

9. Chen S, Zhou Y, Chen Y, Gu J. fastp: an ultra-fast all-in-one FASTQ preprocessor. Bioinformatics. 2018 ;34(17):i884–90.

10. Chin CH, Chen SH, Wu HH, Ho CW, Ko MT, Lin CY. cytoHubba: identifying hub objects and sub-networks from complex interactome. BMC Syst. Biol. 2014; 8(4):1–7. https://doi.org/10.1186/1752-0509-8-s4-s11

11. Ciofu O. Pseudomonas aeruginosa chromosomal beta-lactamase in patients with cystic fibrosis and chronic lung infection. Mechanism of antibiotic resistance and target of the humoral immune response. APMIS. Supplementum. 2003:(116):1–47.

12. Conesa A, Götz S. Blast2GO: a comprehensive suite for functional analysis in plant genomics. Int. J. Plant Genomics. 2008;2008.

13. Davies DG, Parsek MR, Pearson JP, Iglewski BH, Costerton JW, Greenberg EP. The involvement of cell-to-cell signals in the development of a bacterial biofilm. Science. 1998;280(5361):295–8. DOI: 10.1126/science.280.5361.295

14. de Souza A, Vora AH, Mehta AD, Moeller K, Moeller C, Willoughby AJ, Godse CS. SilverSol® a Nano-Silver Preparation: A Multidimensional Approach to Advanced Wound Healing. Wound Healing Research: Current Trends and Future Directions. 2021:355–96. DOI: 10.1007/978-981-16-2677-7_12

15. Deforges L, Le Van Thoi J, Soussy CJ, Duval J. Activity of the amoxicillin-clavulanic acid (augmentin) combination on strains of hospital isolates. Pathol Biol. 1985;33(5):301–8.

16. Dulley JR, Grieve PA. A simple technique for eliminating interference by detergents in the Lowry method of protein determination. Anal. Biochem. 1975;64(1):136–41. https://doi.org/10.1016/0003-2697(75)90415-7

17. El-Fouly MZ, Sharaf AM, Shahin AA, El-Bialy HA, Omara AM. Biosynthesis of pyocyanin pigment by Pseudomonas aeruginosa. J. Radiat. Res. Appl. Sci. 2015;8(1):36–48. https://doi.org/10.1016/j.jrras.2014.10.007

18. Feng QL, Wu J, Chen GQ, Cui FZ, Kim TN, Kim JO. A mechanistic study of the antibacterial effect of silver ions on Escherichia coli and Staphylococcus aureus. J. Biomed. Mater. Res. 2000; 52(4):662–8. https://doi.org/10.1002/1097-4636

19. Gajera G, Godse C, DeSouza A, Mehta D, Kothari V. Potent anthelmintic activity of a colloidal nano-silver formulation (Silversol^®^) against the model worm Caenorhabditis elegans. 2023. https://doi.org/10.21203/rs.3.rs-2091038/v1

20. Gupta P, Mankere B, Chekkoora Keloth S, Tuteja U, Pandey P, Chelvam KT. Increased antibiotic resistance exhibited by the biofilm of Vibrio cholerae O139. J. Antimicrob. Chemother. 2018;73(7):1841–7. https://doi.org/10.1093/jac/dky127

21. Hande S, Sonkar V, Bhoj P, Togre N, Goswami K, Dash D. The role of oxidative and nitrosative stress of silver nanoparticles in human parasitic helminth Brugia malayi: a mechanistic insight. Acta Parasitol. 2021:1–0. https://doi.org/10.1007/s11686-021-00394-4

22. Hassan D, Farghali M, Eldeek H, Gaber M, Elossily N, Ismail T. Antiprotozoal activity of silver nanoparticles against Cryptosporidium parvum oocysts: New insights on their feasibility as a water disinfectant. J. Microbiol. Methods. 2019;165:105698. https://doi.org/10.1016/j.mimet.2019.105698

23. Heales SJ, Bolaños JP, Stewart VC, Brookes PS, Land JM, Clark JB. Nitric oxide, mitochondria and neurological disease. Biochim. Biophys. Acta - Bioenerg. 1999;1410(2):215–28. https://doi.org/10.1016/S0005-2728(98)00168-6

24. Hirshfield I, Barua S, Basu P. Overview of biofilms and some key methods for their study. In: (Goldman E, Green LH, eds) Practical Handbook of Microbiology. 2nd ed. Chapter 42, p. 675–688. Boca Raton: CRC Press; 2009.

25. Hu Y, Wang J, Sun H, Wang S, Liao X, Wang J, An T. Roles of extracellular polymeric substances in the bactericidal effect of nanoscale zero-valent iron: trade-offs between physical disruption and oxidative damage. Env. Sci. Nano. 2019;6(7):2061–73.

26. Jiang S, Wang F, Cao X, Slater B, Wang R, Sun H, Wang H, Shen X, Yao Z. Novel application of ion exchange membranes for preparing effective silver and copper based antibacterial membranes. Chemosphere. 2022;287:132131. https://doi.org/10.1016/j.chemosphere.2021.132131

27. Joshi C, Patel P, Kothari V. Anti-infective potential of hydroalcoholic extract of Punica granatum peel against gram-negative bacterial pathogens. F1000Research. 2019;8(70):70. https://doi.org/10.12688/f1000research.17430.2

28. Joshi C, Patel P, Palep H, Kothari V. Validation of the anti-infective potential of a polyherbal ‘Panchvalkal’ preparation, and elucidation of the molecular basis underlining its efficacy against Pseudomonas aeruginosa. BMC Complementary Altern. Med.. 2019;19(1):1–5. https://doi.org/10.1186/s12906-019-2428-5

29. Kang D, Kirienko DR, Webster P, Fisher AL, Kirienko NV. Pyoverdine, a siderophore from Pseudomonas aeruginosa, translocates into C. elegans, removes iron, and activates a distinct host response. Virulence. 2018;9(1):804–17. https://doi.org/10.1080/21505594.2018.1449508

30. Khan T, Ullah H, Nasar A, Ullah M. Antibiotic Resistance and Sensitivity Pattern of Pseudomonas aeruginosa Obtained from Clinical Samples. Lett. Appl. NanoBioScience. 2023;12(4):112. https://doi.org/10.33263/LIANBS124.112

31. Lacy MK, Klutman NE, Horvat RT, Zapantis A. Antibiograms: new NCCLS guidelines, development, and clinical application. Hosp. Pharm. 2004;39(6):542–53. https://doi.org/10.1177/001857870403900608

32. Langmead B, Salzberg SL. Fast gapped-read alignment with Bowtie 2. Nat. Methods. 2012;9(4):357–9.

33. Lee YH, Cheng FY, Chiu HW, Tsai JC, Fang CY, Chen CW, Wang YJ. Cytotoxicity, oxidative stress, apoptosis and the autophagic effects of silver nanoparticles in mouse embryonic fibroblasts. Biomaterials. 2014; 35(16):4706–15. https://doi.org/10.1016/j.biomaterials.2014.02.021

34. Li Q, Yan W, Yang K, Wen Y, Tang JL. Xanthan gum production by Xanthomonas campestris pv. campestris 8004 using cassava starch as carbon source. Afr. J. Biotechnol. 2012;11(73):13809–13. DOI: 10.5897/AJB11.3774

35. Li WR, Xie XB, Shi QS, Zeng HY, Ou-Yang YS, Chen YB. Antibacterial activity and mechanism of silver nanoparticles on Escherichia coli. Applied microbiology and biotechnology. 2010; 85:1115–22. https://doi.org/10.1007/s00253-009-2159-5

36. Liao Y, Smyth GK, Shi W. featureCounts: an efficient general purpose program for assigning sequence reads to genomic features. Bioinformatics. 2014;30(7):923–30.

37. Litwin CM, Calderwood S. Role of iron in regulation of virulence genes. Clin. Microbiol. Rev. 1993;6(2):137–49. https://doi.org/10.1128/cmr.6.2.137

38. Liu B, Qian SB. Translational reprogramming in cellular stress response. Wiley Interdiscip. Rev.: RNA. 2014;5(3):301–5. https://doi.org/10.1002/wrna.1212

39. Lowry OH, Rosebrough NJ, Farr AL, Randall RJ. Protein measurement with the Folin phenol reagent. J. Biol. Chem. 1951;193:265–75.

40. Matras E, Gorczyca A, Przemieniecki SW, Oćwieja M. Surface properties-dependent antifungal activity of silver nanoparticles. Sci. Rep. 2022;12(1):18046. doi: 10.1038/s41598-022-22659-2

41. Mazur P, Skiba-Kurek I, Mrowiec P, Karczewska E, Drożdż R. Synergistic ROS-associated antimicrobial activity of silver nanoparticles and gentamicin against staphylococcus epidermidis. Int. J. Nanomed. 2020:3551-62. DOI: 10.2147/IJN.S246484

42. McQuillan JS, Shaw AM. Differential gene regulation in the Ag nanoparticle and Ag+- induced silver stress response in Escherichia coli: A full transcriptomic profile. Nanotoxicology. 2014;8(sup1):177–84. https://doi.org/10.3109/17435390.2013.870243

43. Mishra M, Tiwari S, Gomes AV. Protein purification and analysis: next generation Western blotting techniques. Expert Rev. Proteomics. 2017;14(11):1037–53. https://doi.org/10.1080/14789450.2017.1388167

44. Misko TP, Schilling RJ, Salvemini D, Moore WM, Currie MG. A fluorometric assay for the measurement of nitrite in biological samples. Anal. Biochem. 1993;214(1):11–6. https://doi.org/10.1006/abio.1993.144

45. Mohammed M, Mekala LP, Chintalapati S, Chintalapati VR. New insights into aniline toxicity: aniline exposure triggers envelope stress and extracellular polymeric substance formation in Rubrivivax benzoatilyticus JA2. J. Hazard. Mater. 2020;385:121571. https://doi.org/10.1016/j.jhazmat.2019.121571

46. Morales DK, Grahl N, Okegbe C, Dietrich LE, Jacobs NJ, Hogan DA. Control of Candida albicans metabolism and biofilm formation by Pseudomonas aeruginosa phenazines. MBio. 2013;4(1):e00526–12. https://doi.org/10.1128/mbio.00526-12

47. Morones-Ramirez JR, Winkler JA, Spina CS, Collins JJ. Silver enhances antibiotic activity against gram-negative bacteria. Sci. Transl. Med. 2013; 5(190):190ra81. DOI: 10.1126/scitranslmed.3006276

48. O’Toole G, Kaplan HB, Kolter R. Biofilm formation as microbial development. Annu. Rev. Microbiol. 2000;54(1):49–79. https://doi.org/10.1146/annurev.micro.54.1.49

49. Patel P, Joshi C, Kothari V. Antipathogenic potential of a polyherbal wound-care formulation (herboheal) against certain wound-infective gram-negative bacteria. Adv. Pharmacol. Sci. 2019;2019. https://doi.org/10.1155/2019/1739868

50. Pfaller MA, Sheehan DJ, Rex JH. Determination of fungicidal activities against yeasts and molds: lessons learned from bactericidal testing and the need for standardization. Clin. Microbiol. Rev. 2004;17(2):268–80. https://doi.org/10.1128/cmr.17.2.268-280.2004

51. Poole RK. Nitric oxide and nitrosative stress tolerance in bacteria. Biochem. Soc. Trans. 2005;33(1):176–80. https://doi.org/10.1042/BST0330176

52. Qais FA, Shafiq A, Ahmad I, Husain FM, Khan RA, Hassan I. Green synthesis of silver nanoparticles using Carum copticum: Assessment of its quorum sensing and biofilm inhibitory potential against gram negative bacterial pathogens. Microb. Pathog. 2020; 144:104172. https://doi.org/10.1016/j.micpath.2020.104172

53. Ramadan MA, Tawfik AF, Shibl AM, Gemmell CG. Post-antibiotic effect of azithromycin and erythromycin on streptococcal susceptibility to phagocytosis. J. Med. Microbiol. 1995;42(5):362–6. https://doi.org/10.1099/00222615-42-5-362

54. Reeves CM, Magallon J, Rocha K, Tran T, Phan K, Vu P, Yi Y, Oakley-Havens CL, Cedano J, Jimenez V, Ramirez MS. Aminoglycoside 6′-N-acetyltransferase Type Ib [AAC (6′)-Ib]-Mediated Aminoglycoside Resistance: Phenotypic Conversion to Susceptibility by Silver Ions. Antibiotics. 2020;10(1):29. https://doi.org/10.3390/antibiotics10010029

55. Robinson MD, McCarthy DJ, Smyth GK. edgeR: a Bioconductor package for differential expression analysis of digital gene expression data. Bioinformatics. 2010;26(1):139–40.

56. Ruparel FJ, Shah SK, Patel JH, Thakkar NR, Gajera GN, Kothari VO. Network analysis for identifying potential anti-virulence targets from whole transcriptome of Pseudomonas aeruginosa and Staphylococcus aureus exposed to certain anti-pathogenic polyherbal formulations. Drug Target Insights. 2023;17:58. doi: 10.33393/dti.2022.2595

57. Salleh A, Naomi R, Utami ND, Mohammad AW, Mahmoudi E, Mustafa N, Fauzi MB. The potential of silver nanoparticles for antiviral and antibacterial applications: A mechanism of action. Nanomaterials. 2020;10(8):1566. https://doi.org/10.3390/nano10081566

58. Schreiber K, Boes N, Eschbach M, Jaensch L, Wehland J, Bjarnsholt T, Givskov M, Hentzer M, Schobert M. Anaerobic survival of Pseudomonas aeruginosa by pyruvate fermentation requires an Usp-type stress protein. J. Bacteriol. 2006;188(2):659–68. https://doi.org/10.1128/jb.188.2.659-668.2006

59. Shannon P, Markiel A, Ozier O, Baliga NS, Wang JT, Ramage D, Amin N, Schwikowski B, Ideker T. Cytoscape: a software environment for integrated models of biomolecular interaction networks. Genome Res. 2003;13(11):2498–504. doi:10.1101/gr.1239303

60. Singh VK, Jayaswal RK, Wilkinson BJ. Cell wall-active antibiotic induced proteins of Staphylococcus aureus identified using a proteomic approach. FEMS Microbiol. Lett. 2001;199(1):79–84. https://doi.org/10.1111/j.1574-6968.2001.tb10654.x

61. Šinko G, Vinković Vrček I, Goessler W, Leitinger G, Dijanošić A, Miljanić S. Alteration of cholinesterase activity as possible mechanism of silver nanoparticle toxicity. Environ. Sci. Pollut. Res. 2014; 21:1391–400. https://doi.org/10.1007/s11356-013-2016-z

62. Subhadra B, Kim DH, Woo K, Surendran S, Choi CH. Control of biofilm formation in healthcare: Recent advances exploiting quorum-sensing interference strategies and multidrug efflux pump inhibitors. Materials. 2018; 11(9):1676. https://doi.org/10.3390/ma11091676

63. Suzuki J, Kunimoto T, Hori M. Effects of kanamycin on protein synthesis: inhibition of elongation of peptide chains. J. Antibiot. 1970;23(2):99–101. https://doi.org/10.7164/antibiotics.23.99

64. Szklarczyk D, Gable AL, Lyon D, Junge A, Wyder S, Huerta-Cepas J, Simonovic M, Doncheva NT, Morris JH, Bork P, Jensen LJ. STRING v11: protein–protein association networks with increased coverage, supporting functional discovery in genome-wide experimental datasets. Nucleic Acids Res. 2019;47(D1):D607–13. https://doi.org/10.1093/nar/gky1131

65. Tang S, Zheng J. Antibacterial activity of silver nanoparticles: structural effects. Adv. Healthcare Mater. 2018;7(13):1701503. https://doi.org/10.1002/adhm.201701503

66. Trafny EA, Lewandowski R, Zawistowska-Marciniak I, Stępińska M. Use of MTT assay for determination of the biofilm formation capacity of microorganisms in metalworking fluids. World J Microbiol Biotechnol. 2013; 29(9):1635–43. https://doi.org/10.1007/s11274-013-1326-0

67. Turnidge JD. Susceptibility test methods: general considerations. Manual of clinical microbiology. 2015:1246–52.

68. Unni KN, Priji P, Geoffroy VA, Doble M, Benjamin S. Pseudomonas aeruginosa BUP2—A novel strain isolated from malabari goat produces Type 2 pyoverdine. Adv. Biosci. Biotechnol. 2014;5(11):874. DOI: 10.4236/abb.2014.511102

69. Untergasser A, Nijveen H, Rao X, Bisseling T, Geurts R, Leunissen JA. Primer3Plus, an enhanced web interface to Primer3. Nucleic Acids Res. 2007;35(suppl_2):W71–4.

70. Wahab MA, Luming L, Matin MA, Karim MR, Aijaz MO, Alharbi HF, Abdala A, Haque R. Silver micro-nanoparticle-based nanoarchitectures: Synthesis routes, biomedical applications, and mechanisms of action. Polymers. 2021;13(17):2870. https://doi.org/10.3390/polym13172870

71. Weiner LM, Webb AK, Limbago B, Dudeck MA, Patel J, Kallen AJ, Edwards JR, Sievert DM. Antimicrobial-resistant pathogens associated with healthcare-associated infections: summary of data reported to the National Healthcare Safety Network at the Centers for Disease Control and Prevention, 2011–2014. Infect. Control Hosp. Epidemiol. 2016; 37(11):1288–301.

72. Wesselink E, Koekkoek WA, Grefte S, Witkamp RF, Van Zanten AR. Feeding mitochondria: Potential role of nutritional components to improve critical illness convalescence. Clin. Nutr. 2019;38(3):982–95. https://doi.org/10.1016/j.clnu.2018.08.032

73. Yang L, Haagensen JA, Jelsbak L, Johansen HK, Sternberg C, Høiby N, Molin S. In situ growth rates and biofilm development of Pseudomonas aeruginosa populations in chronic lung infections. J. Bacteriol. 2008; 193(9), 2760–2768. https://doi.org/10.1128/jb.01581-07

74. Ye J, Zhang Y, Cui H, Liu J, Wu Y, Cheng Y, Xu H, Huang X, Li S, Zhou A, Zhang X. WEGO 2.0: a web tool for analyzing and plotting GO annotations, 2018 update. Nucleic Acids Res. 2018;46(W1):W71–5.

75. Yin IX, Zhang J, Zhao IS, Mei ML, Li Q, Chu CH. The antibacterial mechanism of silver nanoparticles and its application in dentistry. Int. J. Nanomed. 2020;17:2555–62. DOI: 10.2147/IJN.S246764

76. Yin IX, Zhang J, Zhao IS, Mei ML, Li Q, Chu CH. The antibacterial mechanism of silver nanoparticles and its application in dentistry. Int. J. Nanomed. 2020: 2555–62. DOI: 10.2147/IJN.S246764

77. Zhang XF, Liu ZG, Shen W, Gurunathan S. Silver nanoparticles: synthesis, characterization, properties, applications, and therapeutic approaches. Int. J. Mol. Sci. 2016;17(9):1534. https://doi.org/10.3390/ijms17091534

