## Supplementary File for "Identifying molecular targets of a colloidal nanosilver formulation (Silversol^®^) in multidrug resistant *Pseudomonas aeruginosa*"

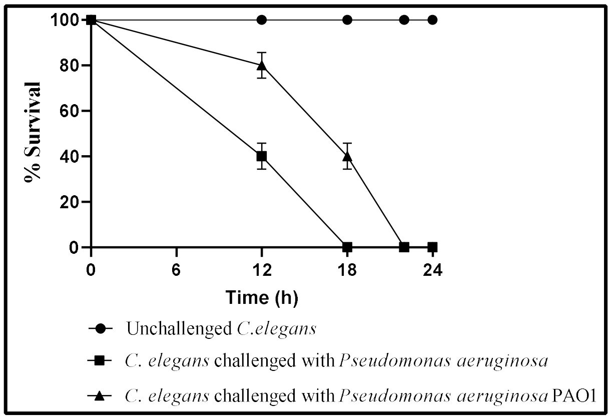


**Figure S1. MDR strain of *P. aeruginosa* used in our study could kill the host worms faster than the PAO1 strain**

When worms in liquid media were challenged with either strain of *P. aeruginosa,* it took 18 h for the MDR strain to kill cent percent of the worm population as against 22 h required by PAO1 to achieve same effect. At the 12-hour and 18-hour end-points, MDR could kill 40%±1*** and 40%±5.7*** more worms than PAO1. ***$p<0.001$

**Table S1. Comparison of antibiograms of *Pseudomonas aeruginosa* (MDR strain used in this study) and PAO1 generated through Kirby-Bauer Disc Diffusion assay**

| Sr. No. | Antibiotic | Concentration (µg per disc) | *Pseudomonas aeruginosa* (MDR) | *Pseudomonas aeruginosa* PAO1 |
|---|---|---|---|---|
| 1 | Ceftriaxone | 30 | S | S |
| 2 | Colistin | 10 | S | S |
| 3 | Ciprofloxacin | 5 | S | S |
| 4 | Co-Trimoxazole | 25 | R | R |
| 5 | Imipenem | 10 | S | S |
| 6 | Ticarcillin | 75 | S | S |
| 7 | Streptomycin | 25 | S | S |
| 8 | Sparfloxacin | 5 | S | S |
| 9 | Cefpodoxime | 10 | S | S |
| 10 | Nalidixic Acid | 30 | S | S |
| 11 | Moxifloxacin | 5 | S | S |
| 12 | Gentamicin | 10 | S | S |
| 13 | Gatifloxacin | 5 | S | S |
| 14 | Ofloxacin | 5 | S | S |
| 15 | Tobramycin | 10 | S | S |
| 16 | Norfloxacin | 10 | S | S |
| 17 | Amikacin | 30 | S | S |
| 18 | Levofloxacin | 5 | S | S |
| 19 | Augmentin | 30 | R | R |
| 20 | Kanamycin | 30 | S | S |
| 21 | Vancomycin | 30 | R | R |
| 22 | Cefixime | 5 | R | R |
| 23 | Tetracycline | 30 | I | S |
| 24 | Rifampicin | 5 | I | R |
| 25 | Clindamycin | 2 | R | R |
| 26 | Chloramphenicol | 30 | R | S |
| 27 | Cefepime | 30 | S | S |
| 28 | Doxycycline Hydrochloride | 30 | I | I |
| 29 | Cefotaxime | 30 | S | S |
| 30 | Nitrofurantoin | 300 | R | R |
| 31 | Ampicillin | 10 | R | R |

Antibiotic susceptibility profile of the organisms was generated using the antibiotic discs- Icosa G-I Minus (HiMedia, Mumbai) through disc diffusion assay on cation-adjusted Mueller-Hinton agar (HiMedia) as per National Committee for Clinical Laboratory Standards (NCCLS) guidelines (https://doi.org/10.1177/001857870403900608). The zones of inhibition were measured and the interpretation (S - sensitive, I - intermediate, R - resistant) was drawn as per zone size interpretative chart provided by the manufacturer.

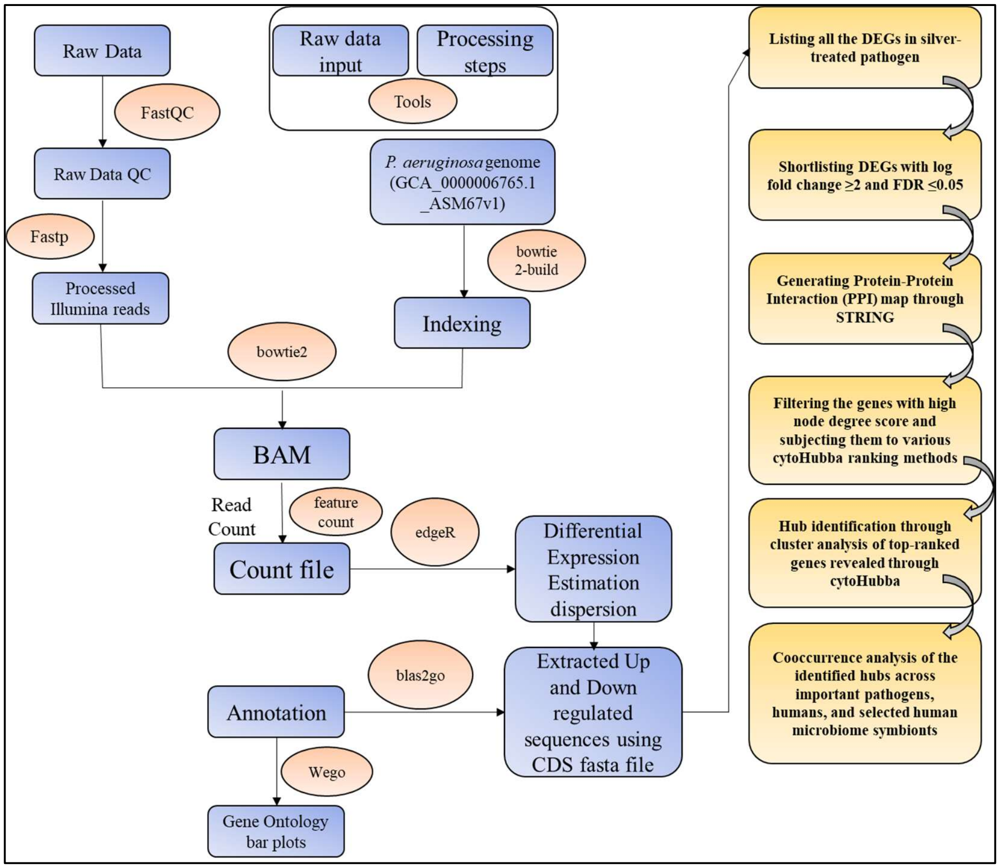


**Figure S2: A schematic presentation of the methodology/workflow employed for whole transcriptome and network analysis**

DEG: Differentially Expressed Genes

**Table S2. Quantification of extracted RNA**

| Sr. No. | Sample Name | Volume (µL) | ng/ µL | $OD_{260}/OD_{280}$ | $OD_{260}/OD_{230}$ | Quantity (ng/ µL) | RIN value |
|---|---|---|---|---|---|---|---|
| 1 | Control | 17 | 2101.9 | 2.15 | 2.47 | 1440 | 6.5 |
| 2 | Experimental | 17 | 881.6 | 2.14 | 2.95 | 800 | 8.3 |

**Table S3. Library quantification and insert size analysis**

| Sr. No. | Sample Name | ng/ µL | Insert size | Index |
|---|---|---|---|---|
| 1 | Control | 21 | 149,324,439 | 22 |
| 2 | Experimental | 24.4 | 366,439 | 23 |

**Table S4. Temperature profile for RT-PCR assay**

| Temperature (°C) | Time (s) | Remarks |
|---|---|---|
| PCR cycles (45 cycles) | | |
| 95 | 15 | Denaturation temperature |
| 59 | 60 | Annealing temperature |
| **Melt curve stage** | | |
| 95 | 15 | |
| 60 | 60 | |
| 95 | 15 | |

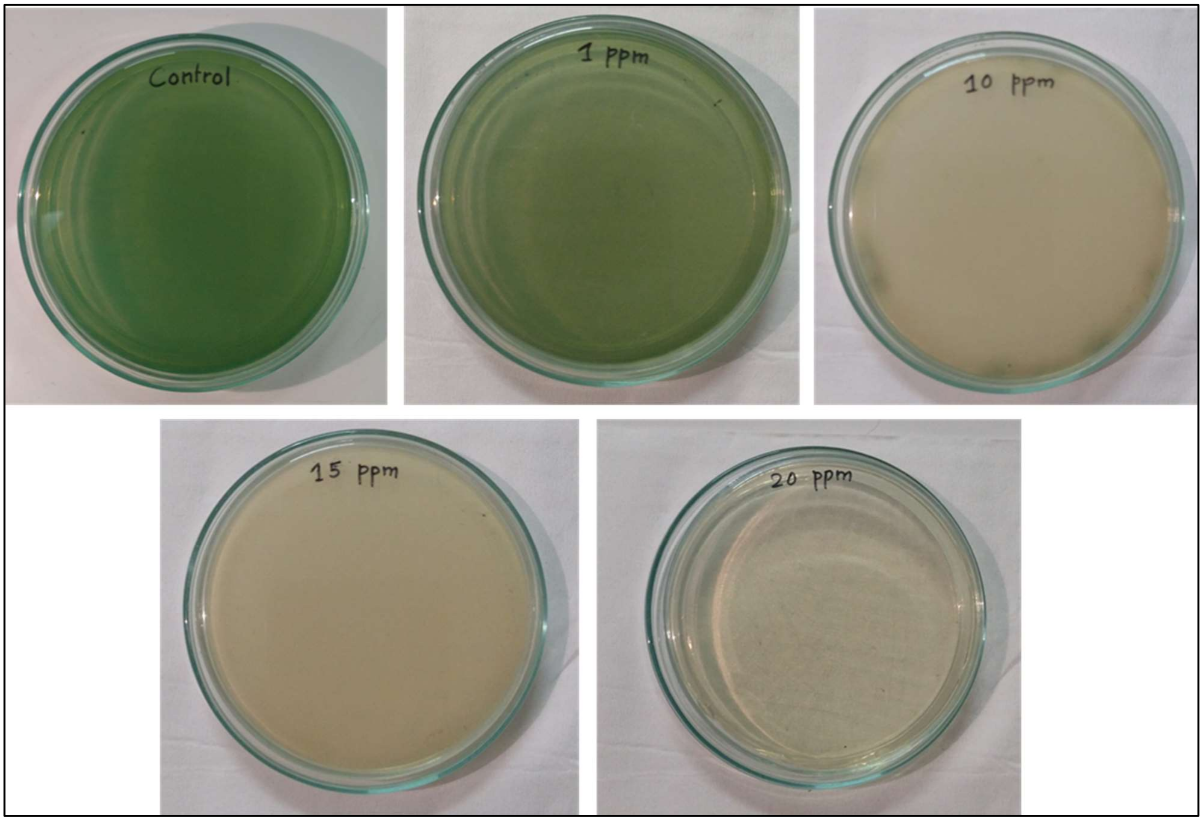


**Figure S3. MBC of Silversol® against *P. aeruginosa* was between 16-20 ppm**

Cells grown in presence of Silversol® were subsequently plated onto Pseudomonas agar. Those coming from 10-15 ppm silver tubes displayed non-pigmented growth, while those from 20-ppm failed to give rise to any visible growth.

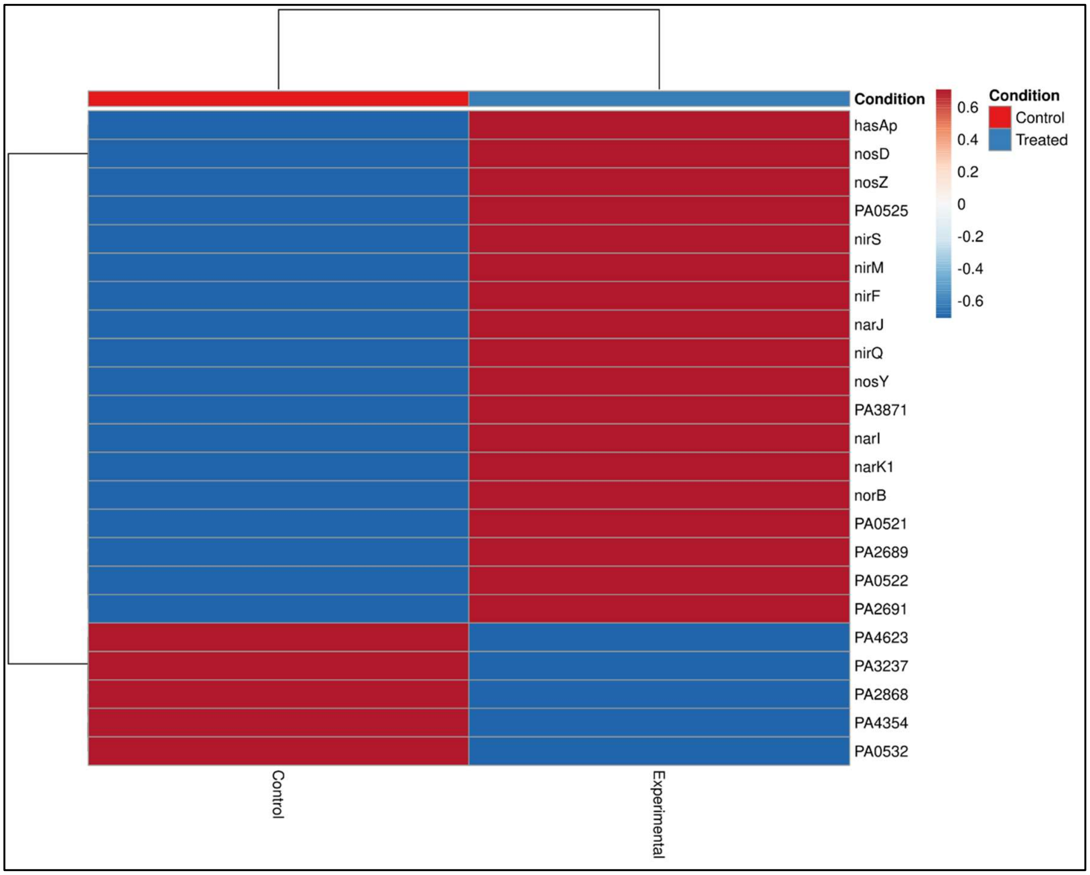


**Figure S4. Heat map of DEGs in Silversol®-exposed *P. aeruginosa***

Heat map generated using the online software tool ClustVis (https://biit.cs.ut.ee/clust vis/) showing upregulated and downregulated genes with FDR<0.05 and log fold change '± 2'

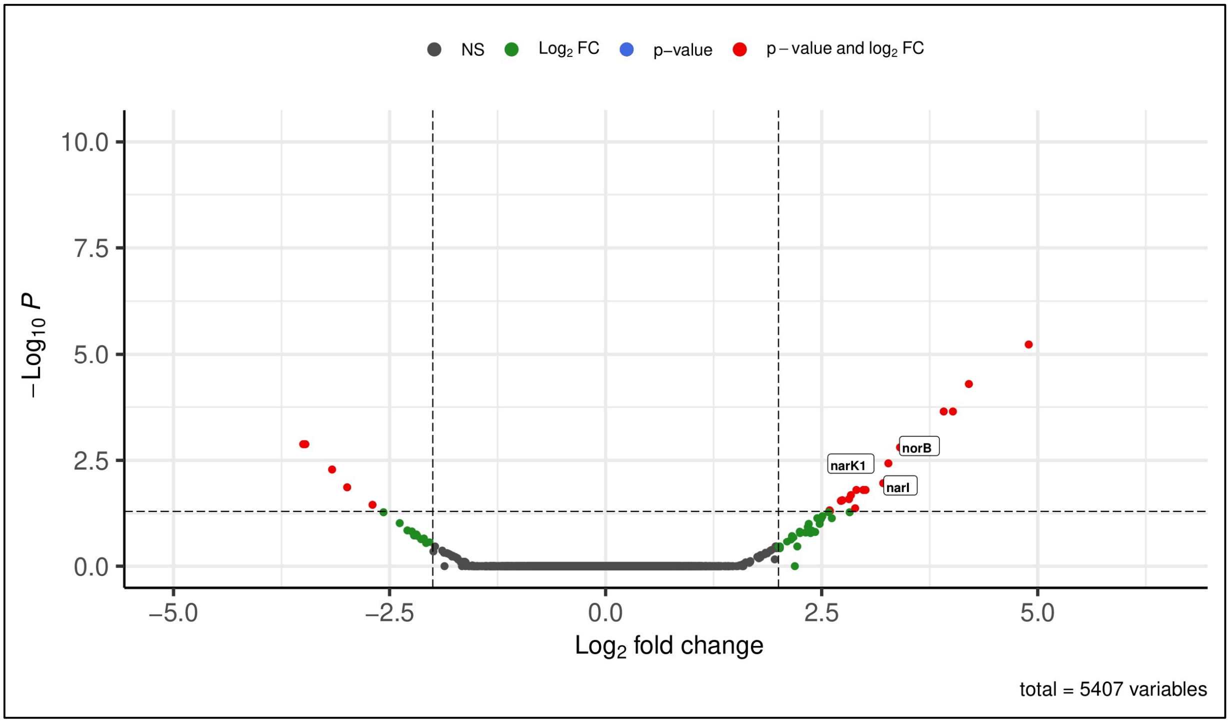


**Figure S5. Volcano Plot of experimental versus control samples**

Volcano plot of expressed genes of experimental culture compared to control culture. The y-axis illustrates −log 10 p values, and the x-axis corresponds to a log 2-fold change of gene expression between both cultures. The red points represent differently expressed genes satisfying the dual criteria of FDR< 0.05 and log fold change ≥2.
